# Distinct fibroblast and perivascular senotypes define spatial niches that regulate fibrosis

**DOI:** 10.64898/2026.06.08.730636

**Authors:** Alexandra N. Rindone, Sushma Nagaraj, Anna Cho, Maria Browne, Kavita Krishnan, Ricky S. Adkins, Daniel Lesperance, Joshua Orvis, Prarthana Sanjay Daswani, Joscelyn C. Mejías, Jacob P. Rose, Christina D. King, Anna Ruta, Frank Haoning Yu, Kofi O. Boahene, Birgit Schilling, Anup A. Mahurkar, Owen R. White, Elana J. Fertig, Jennifer H. Elisseeff

**Affiliations:** Translational Therapeutics & Regenerative Engineering Center, Department Chemical and Biomolecular Engineering and Biomedical Engineering, Johns Hopkins University, Baltimore, MD; Institute for Genome Sciences, University of Maryland School of Medicine, Baltimore, MD; Wallace H. Coulter Department of Biomedical Engineering, Georgia Tech and Emory University, Atlanta, GA; Buck Institute for Research on Aging, Novato, CA; Department of Otolaryngology – Head and Neck Surgery, Johns Hopkins University School of Medicine, Baltimore, MD; Greenebaum Comprehensive Cancer Center, Department of Medicine and Department of Epidemiology, University of Maryland School of Medicine, Baltimore, MD

## Abstract

Fibrotic conditions contribute to significant global morbidity and mortality. Yet the underlying processes that orchestrate fibrosis remain poorly understood due to the cellular and spatial complexity of the stromal, immune, and vascular compartments that regulate fibrotic disease progression. Senescent cells (SnCs) have been implicated in fibrosis, but their roles are unclear, as evidence indicates that they serve both pathogenic and reparative functions. Here, we show that fibrosis-associated SnCs contain functionally divergent senotypes that are organized into distinct spatial niches. Using integrated single-cell and spatial transcriptomics analyses and hierarchical factorization in a murine fibrosis model, we identify fibroblast and perivascular SnC subpopulations that upregulate diverse programs related to extracellular matrix (ECM) production, immune signaling, and vascular remodeling. Fibroblast senotypes localize to discrete microenvironments with distinct tissue architectures, including niches associated with fibrotic signaling, immune activity, and cartilage development. Perivascular SnCs occupy interfaces between fibrotic signaling and immune-active niches and upregulate vascular and fibrotic remodeling pathways. Depletion of pericyte-lineage SnCs increases vascular maturation and fibrotic ECM deposition, providing mechanistic validation of the beneficial role these SnCs play in vascular remodeling and fibrosis modulation. In addition, using a new web-based infrastructure to query our senotype gene signatures in public datasets, we demonstrate that these senotypes are conserved across different murine and human fibrotic conditions. These findings establish senescence as a spatially organized regulator of fibrosis and identify perivascular senescence as a link between vascular remodeling and fibrotic outcomes.

## Introduction

Fibrosis is a major contributor to global morbidity and mortality and can arise in many tissue types in response to stimuli such as injury, chronic inflammation, and aging. Despite shared hallmarks, including extracellular (ECM) deposition and fibroblast activation, fibrotic diseases remain largely resistant to therapy and the underlying mechanisms governing fibrosis are not yet fully understood. Recent studies indicate that fibrosis is not simply the result of persistent fibroblast activation, but rather emerges from dysregulated tissue repair processes shaped by interactions between stromal, immune, and vascular compartments^1–5^.

Cellular senescence has emerged as a central feature of aging and fibrotic diseases. Senescent cells (SnCs) accumulate in fibrotic tissues across multiple organs^6–8^ and secrete bioactive factors collectively termed as the senescence-associated secretory phenotype (SASP), which can influence inflammation, ECM remodeling, and tissue function. While senescence has been broadly implicated in fibrosis, its role remains paradoxical: SnC depletion can reduce fibrosis in some contexts^6–9^ but impair tissue repair or exacerbate fibrotic pathology in others^10–14^. These divergent outcomes suggest that senescence is not a uniform state and instead consists of functionally distinct phenotypes (senotypes) that depend on cellular identity, microenvironment, and disease stage.

A critical gap in the field is our lack of understanding of how senotypes are organized within tissue and how they interact with their local microenvironments to regulate fibrotic outcomes. In particular, fibrosis is increasingly recognized as a spatially structured process, where distinct cellular niches, including immune-rich, fibrotic, and vascular regions, regulate inflammation, ECM production, and tissue remodeling^1–3,15,16^. Recent advances in bioinformatics methods for intercellular communication and spatial molecular profiling have enabled us to delineate the unexplored contribution of SnCs to these niche-level processes. Furthermore, while fibroblasts have been the primary focus of fibrosis research, these computational assays can explore the role of senescent perivascular cells and vascular remodeling in shaping fibrosis. These cells have received comparatively little attention, despite evidence that pericytes and vascular-associated cells regulate both tissue repair and fibrotic progression^4,5,17,18^.

Here, we hypothesize that senescence contributes to fibrosis through distinct, spatially organized senotypes embedded within different stromal, immune, and vascular niches, rather than through a uniform accumulation of SnCs. To test this hypothesis, we apply a multimodal platform integrating single-cell transcriptomics and spatial transcriptomics in a murine model of fibrosis with robust senescence induction. Using transfer learning and hierarchical factorization, we identify functionally distinct fibroblast and perivascular senotypes and define their niche-specific transcriptional programs related to fibrosis, immune signaling, and tissue and vascular remodeling. We further functionally validate that perivascular senescence regulates vascular maturation and modulates fibrosis progression using a new pericyte-lineage SnC elimination model. Finally, leveraging a web-based infrastructure, gEAR^19^, we show that these fibroblast and perivascular cell senotypes are present in murine models of muscle regeneration and bleomycin-induced pulmonary fibrosis, as well as in human keloid, illustrating the conservation of such senotypes across different biological conditions and species.

Through integrated experimental and computational approaches, our study suggests that senotypes are not merely heterogeneous cellular states but components of spatially organized niche ecosystems that shape tissue structure and function. Distinct senotypes were associated with unique cellular compositions, extracellular matrix architectures, and immune microenvironments, indicating that senescent cells participate in the construction of higher-order tissue organization. These findings support a hierarchical model in which senescence gives rise to specialized senotypes, senotypes establish local niches through SASP-mediated interactions, and interactions among niches ultimately govern tissue remodeling and fibrosis.

### Multiple fibroblast and perivascular subpopulations are senescent in murine fibrosis

We used a murine model that reproducibly induces chronic inflammation, fibrosis, and senescence to study heterogeneous senotypes associated with processes involved in fibrosis. This model implants polycaprolactone (PCL) particles into a volumetric muscle loss injury to induce a foreign body response that causes sustained inflammation and fibrosis around implant particles. This response results in an accumulation of p16+ and p21+ SnCs (5-15% of total cells per marker) that provides a reliable model to study heterogeneous senotypes *in vivo* (**Ext. Data 1A-D**). Previously, we generated single cell RNA-sequencing (scRNA-seq) data from this model^20–22^ to study the interactions between T cell subpopulations and stromal/vascular cells in fibrosis. Here, we leveraged these same data to conduct an analysis of the senotypes associated with fibrosis. This dataset contains 18,201 cells divided into stromal, vascular, and immune cell clusters, including fibroblasts (*Col1a1, Dcn*), endothelial cells (ECs) (*Cdh5, Pecam1*), lymphatic ECs (*Lyve1, Prox1*), perivascular cells (*Rgs5, Acta2*), myeloid cells (*Itgam, Itgax, Cd68*), and T/NK cells (*Cd3e, Nkg7*) (**Fig. 1A**). The fibroblast subclusters include two progenitor populations (*Pi16+, Bmp5+*), activated fibroblasts, *Lrrc15*+ myofibroblasts, and cartilage-like fibroblasts, as previously reported^21,22^. In addition, to extend our analyses to the myeloid compartment, we conducted subclustering to identify the different subtypes of myeloid cells present in the fibrosis tissue. Through this analysis, we identified subclusters corresponding with Neutrophils, Monocytes/Early Macrophages, *Spp1*^hi^ Macrophages, and *Mrc1*+ Macrophages (**Supplemental Data 1**).

**Figure 1.**
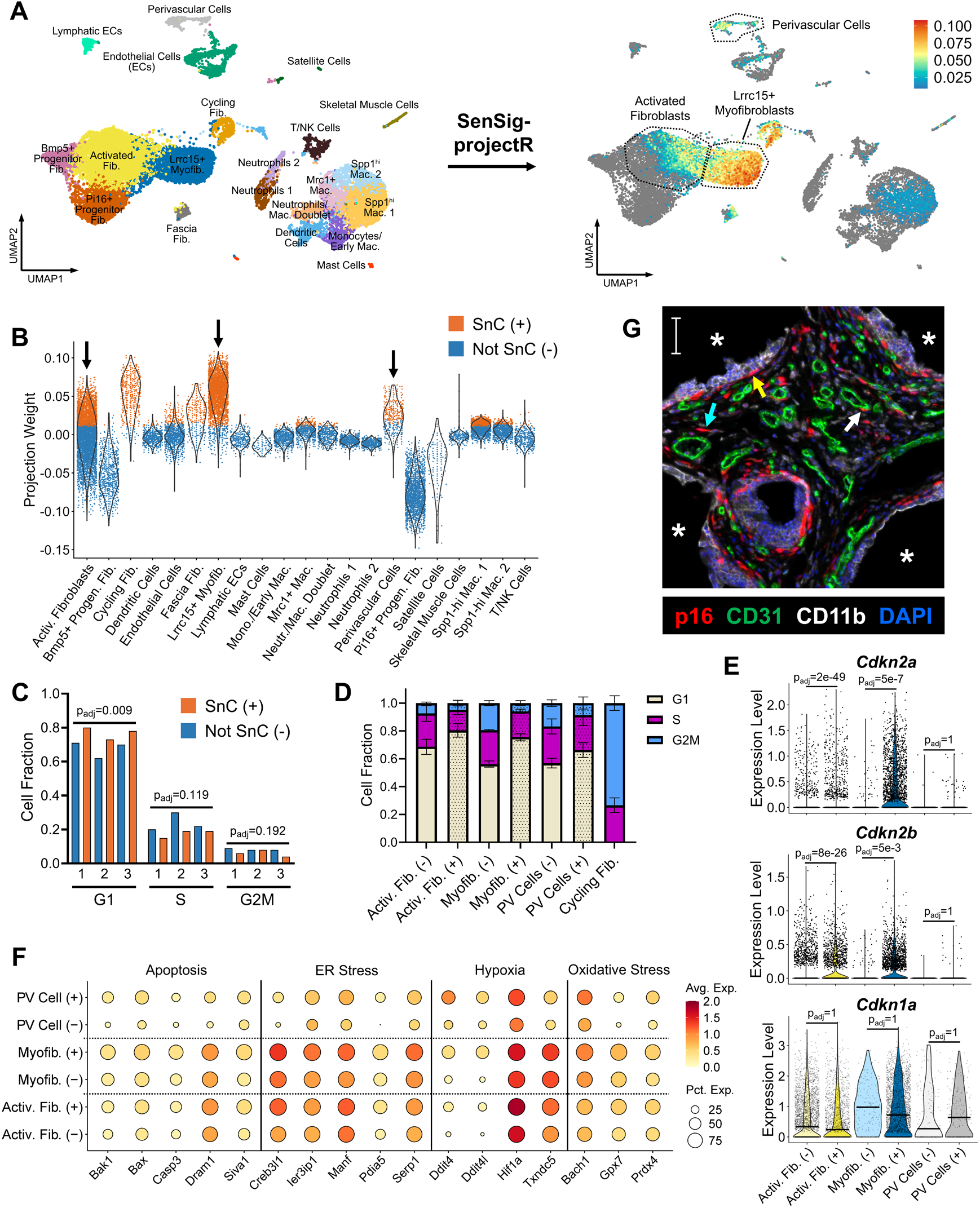
SenSig-projectR identifies fibroblast and perivascular subpopulations enriched in senescent cells (SnCs) in murine fibrosis model. **A)** Application of transfer learning using SenSig to a scRNA-seq dataset of a murine biomaterial fibrosis model. Cells with a positive projection weight and p-value <0.01 were considered senescent. Right UMAP plot scale represents SenSig projection weight in cells classified as SnC. **B)** SenSig projection weights of the cells in each cluster. Activated fibroblasts (Activ. Fib.), *Lrrc15*+ myofibroblasts (Myofib.), perivascular (PV) cells are highlighted with the arrows. **C)** Fraction of SnCs (+) and Not SnCs (-) from activated fibroblast, *Lrrc15*+ myofibroblast, and perivascular cell clusters in G1, S, and G2M phases assigned via Seurat cell cycle scoring. Each number corresponds to a biological replicate (n=3). Cell fractions in SnC and Not SnC groups were compared in each cell cycle phase using a paired t-test with Paired t-test with Holm-Šídák method. **D)** Comparison of Seurat-assigned cell cycle phase by cluster. Cycling fibroblasts are shown in the lined bars as a positive control. Values are mean ± standard deviation. **E)** Normalized expression of cell cycle inhibitors in fibroblast and perivascular SnCs and not SnCs. n=3, Wilcoxon Rank Sum test with Bonferroni correction. **F)** Expression of genes associated with cellular stress and apoptosis in fibroblast and perivascular subpopulations. Dot color represents average normalized expression in each cell subpopulation. Dot size represents percentage of cells within each subpopulation that express the gene. **G)** Immunostaining confirming the presence of senescent p16+ fibroblast subpopulations (yellow and teal) and perivascular cells (white). Scale bar: 50 µm. *PCL particles

Using our annotated single-cell dataset, we applied a transfer learning approach, projectR^23,24^, with our previously published *in vivo* senescence signature, SenSig^25^, to predict which cells were senescent in the fibrotic tissue (**Fig. 1A**). SenSig is an *in vivo*-derived signature of senescence obtained through bulk RNA-seq and differential expression analysis between p16+ and p16-CD45-CD31-CD29+ stromal cells in our fibrosis model. We previously showed that SenSig detects heterogeneous SnC subpopulations in scRNA-seq datasets of different biological conditions, including idiopathic pulmonary fibrosis and basal cell carcinoma^25^. In this study, we apply the transfer learning method projectR^23^ that uses a generalized least squares regression and a Wald test to quantify which cells in our scRNA-seq dataset have gene expression profiles that significantly correlate with the gene weights defined in SenSig. We then classified cells with a positive projection weight (coefficient) from the least squares regression and p-value from the Wald test less than 0.01 as senescent. From this analysis, we found that the most prevalent SnC subpopulations were activated fibroblasts and *Lrrc15+* myofibroblasts (**Fig. 1B**). We also observed a higher percentage of SnCs in the perivascular cell cluster relative to other non-fibroblast cell types—although perivascular SnCs were lower in abundance than fibroblast SnCs (**Fib. 1B**). Additionally, the cycling fibroblast and fascia fibroblast clusters exhibited an elevated SenSig projection weight due to their expression of fibrotic genes (**Fig. 1B**, **Ext. Data 2**). However, we confirmed that these clusters are not senescent based on their expression of proliferative genes (cycling fibroblasts) or their lack of protein expression of p16 and p21 (fascia fibroblasts, **Ext. Data 1B**) and therefore omitted them from further analysis.

To confirm our predictions of senescence, we evaluated gene and protein expression of cell cycle regulators and other senescence-associated genes in the designated subpopulations of fibroblasts and perivascular cells. Compared to non-SnCs, SnCs in activated fibroblasts, myofibroblasts, and perivascular cells had a higher fraction of cells in the Seurat-assigned^26,27^ G1 phase (**Fig. 1D-E**) and higher expression of cell cycle inhibitors *Cdkn2a* (p16) and *Cdkn2b* (p15) (**Fig. 1F**). Furthermore, SnCs expressed higher levels of genes related to apoptosis, ER stress, hypoxia, and oxidative stress, processes that have been linked to senescence (**Fig. 1G**). Notably, all SnC subtypes upregulated *Bak1* and *Bax*—genes that encode for mitochondrial outer membrane permeabilization and have been recently linked to senescence *in vivo*^28^. Protein-based multiplex immunofluorescence staining confirmed the expression of p16 in cells with a fibroblast morphology and in perivascular cells (**Fig. 1C**). By contrast, p16 expression was mostly absent in CD11b+ myeloid cells and CD31+ ECs, confirming that those cells were not the predominant p16+ SnC subtypes (**Fig. 1C, Ext. Data 1E**). We observed minimal protein co-expression of p16 and proliferative marker Ki67, confirming that p16+ cells were consistent with a senescence phenotype (**Ext. Data 1B-D**). Some p16+ cells were also positive for p21—another cell cycle inhibitor linked to senescence. However, the majority of cells were only positive for either p16 or p21, the latter of which was observed in regions consistent with biomaterial-associated myeloid cells (**Ext. Data 1A-C**). Due to the lack of *in vivo*-validated gene signatures for p21+p16-cells, we focused our analysis on cells with positive protein p16 expression, which included fibroblasts and perivascular cells.

### Fibroblast and perivascular senotypes upregulate distinct SASP profiles

To characterize fibroblast and perivascular senotypes, we performed differential gene expression analysis of SnCs versus non-SnCs within each cluster. We identified 2985, 938, and 490 genes that were upregulated in the activated fibroblast SnCs, myofibroblast SnCs, and perivascular SnCs, respectively (**Supplemental Data 2**). As SASP factors are recognized as important phenotypic traits of SnCs, we focused our analysis by annotating SASP factors using published databases of secreted ligands and ECM molecules^29–31^. We excluded genes predominantly associated with cell membrane proteins using the SURFY database^32^, leaving a comprehensive list of SASP factors for analysis (**Supplemental Data 3**). Through this analysis, we identified a total of 272 SASP factors upregulated among the different SnC subtypes (**Fig. 2A**). All SnC subtypes upregulated SASP factors related to ECM formation and remodeling, including collagens (e.g. *Col1a1/2, Col4a1/2, Col5a1/2/3*), ECM glycoproteins and proteoglycans (*Bgn, Mgp, Tnc*), ECM regulators (*Timp1, Serpine2, Serpinh1*), and growth factors (*Bmp1, Pdgfa*) (**Fig. 2B**). We validated protein-level expression of these SASP factors using data-independent acquisition mass spectrometry (DIA-MS)^33–35^ (**Ext. Data 3A**). DIA-MS confirmed the protein expression of 169 of 272 (62.1%) SASP factors (**Ext. Data 3B-C**).

**Figure 2.**
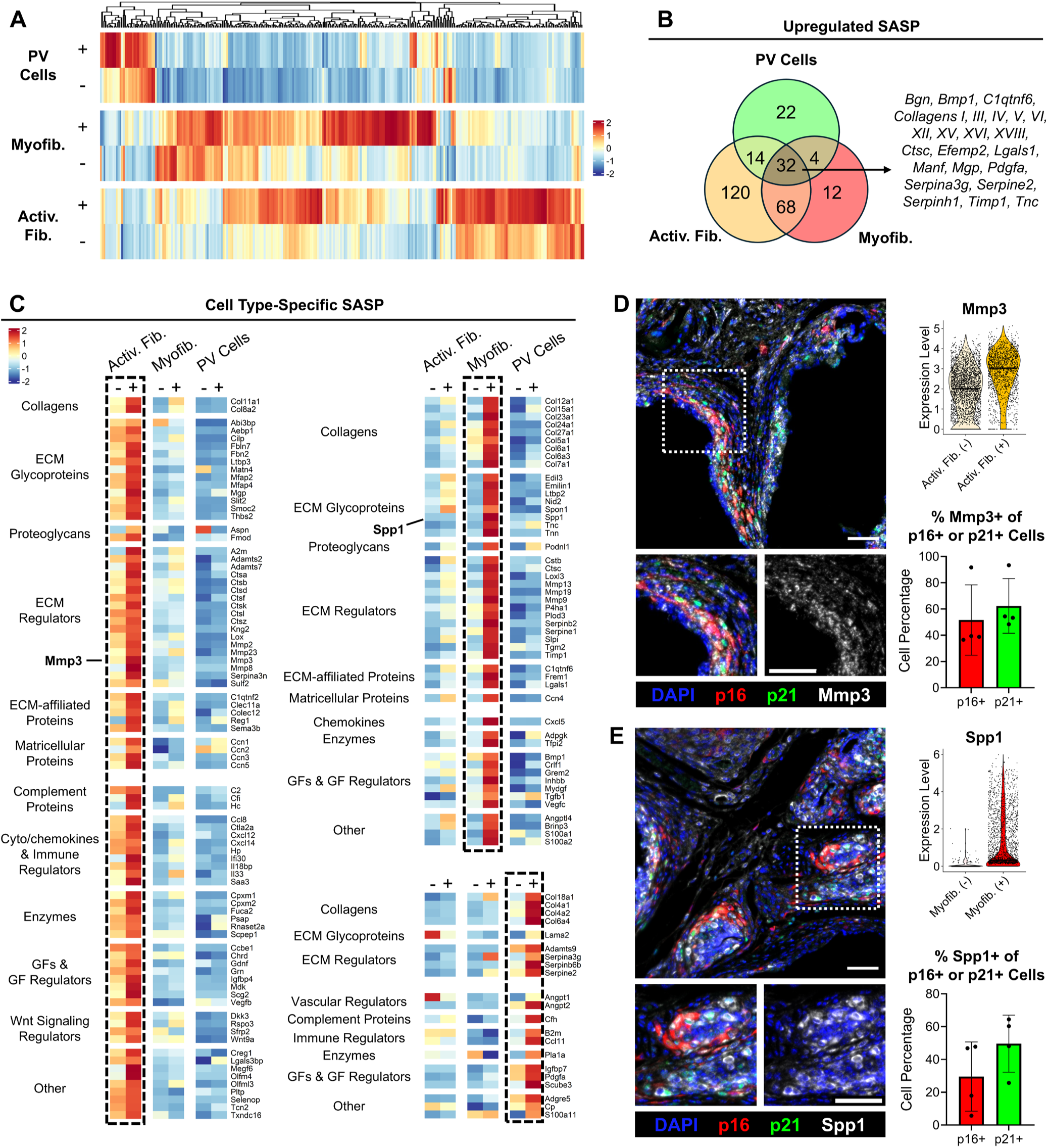
Differential expression analysis predicts senotype-specific SASP. **A)** Heatmap displaying overview of SASP factors upregulated in at least one of the three SnC subtypes. Heatmap colors represent average normalized expression scaled by row. **B)** Venn diagram showing number of SASP genes that are shared and unique to each SnC subtype. **C)** Heatmaps showing cell type-specific SASP in each SnC subtype, derived from **A**. Heatmap colors represent average normalized expression scaled by row. **D-E)** RNA and protein expression of Mmp3 (**D**) and Spp1 (**E**) in SnCs. IF images and bar plots display colocalization of senescence markers, p16 and p21, with Mmp3 and Spp1. Bar plots: Mean ± standard deviation, n=4 biological replicates. Scale bar: 50 μm

While SnCs shared expression of some SASP factors, we found that the levels of expression between the identified SASP factors varied between the SnC subtypes (**Fig. 2C**). Activated fibroblast SnCs expressed SASP genes related to ECM degradation (*Mmp2/3/8*, *Ctsb/k*, *Adamts7*), fibrocartilage matrix formation (*Col11a1, Fmod, Mgp*), inflammation (*Ccl8, Cxcl12, Saa3*), and Wnt signaling (*Wnt9a, Dkk3, Rspo3*). By contrast, myofibroblast SnCs expressed SASP genes related to fibrotic ECM production and remodeling, including several fibrillar and non-fibrillar collagens (*Col5a1, Col6a1/3, Col12a1, Col24a1*), ECM glycoproteins (*Ltbp2, Spp1, Tnc*), and ECM regulators (*Mmp9/13, Serpine1, Timp1*). Perivascular SnCs uniquely expressed SASP genes related to vascular remodeling (*Col4a1/2*, *Col18a1*, *Angpt1/2*, *Adamts9*). We validated the protein expression of Mmp3 and Spp1 in SnCs using multiplex IF with p16 and p21 (**Fig. 2D-E**). We selected these proteins for validation due to their specificity to activated fibroblast and myofibroblasts, respectively, and to the availability of well-validated immunohistochemistry antibodies to these proteins. We identified subsets of p16+ and p21+ cells that co-expressed Mmp3 and Spp1, further validating our transcriptomics predictions that these SASP factors are expressed by some, but not all, SnCs.

To further understand the senotype-specific gene profiles, we compared differentially expressed genes from each SnC subtype with senescence signatures published in other *in vitro* and *in vivo* contexts^25,36–39^ via gene set enrichment analysis (**Ext. Data 4A**). Myofibroblast SnCs had significant positive normalized enrichment scores (NES) for a p16+ cell signature obtained from bleomycin-injured lung^37^, while both fibroblast SnC subtypes had significant negative NES for a gene set downregulated in *in vitro* IR-induced dermal fibroblasts^25^ (**Ext. Data 4B**). By contrast, perivascular SnCs had a positive NES for the *in vitro* p21 PASP^38^ and *in vivo* Day 7 FAP muscle injury^36^ signatures, illustrating their phenotypic differences compared to fibroblast SnCs. All SnC subtypes were negatively enriched for multiple signatures obtained from SPiDER-GAL+ cells in an acute muscle injury^36^. These data replicate a recent finding showing that genes upregulated in p16+p21+ cells were correlated with SenSig but not with the SPiDER-GAL signatures^40^. These trends indicate a distinction between SnCs expressing cell cycle inhibitors versus SA-β-Gal.

We compared cell-based module scores computed with Seurat v5^27^ for select signatures to *Cdkn2a* expression and SenSig projection weight to further investigate these senescence signatures. Through this comparison, we found that SenMayo^39^ was not expressed in similar cell populations as *Cdkn2a*+ and SenSig-high cells in our dataset (**Ext. Data 4C**). By contrast, the lung injury (14 DPI)^37^ and p21 PASP^38^ signatures were more highly expressed in SenSig-high cells in the fibroblast and perivascular cell clusters, respectively (**Ext. Data 4C**). These data show that some senescence signatures are enriched for certain senotypes over others, and their accuracy may be affected when applied to different tissues and physiological states.

### Fibroblast SnCs exhibit distinct senotypes related to fibrosis, immune signaling, and tissue remodeling

Differential expression analysis revealed heterogeneous SASP factors across the fibroblast subclusters, suggesting that traditional clustering methods do not fully capture the complexity of fibroblast senotypes. Whereas clustering methods seek mutually exclusive features, non-negative matrix factorization (NMF) methods are specifically designed to infer gene expression changes that may co-occur between cellular populations (i.e., gene expression patterns) as observed in analysis of SASP factors spanning multiple fibroblast clusters^41^. To better understand the biological patterns expressed in fibroblast senotypes, we applied the Bayesian NMF method CoGAPS^42^ to all of the fibroblast subclusters in the scRNA-seq dataset (**Fig. 3A**). This unsupervised learning method enables the unbiased detection of biological patterns associated with the co-expression of multiple genes that are upregulated by fibroblast senotypes, and it is particularly effective among classes of NMF methods at detecting gene patterns in sparse datasets such as scRNA-seq data^24,42,43^. The primary input parameter to this method is the number of patterns (*k*) sought in the data, which previous studies have shown can be inferred by seeking a drop in the accuracy of the fit of the data quantified through a Chi-squared test^42,44,45^, observed first at *k* = 12 (Low Resolution CoGAPS gene co-expression patterns).

**Figure 3.**
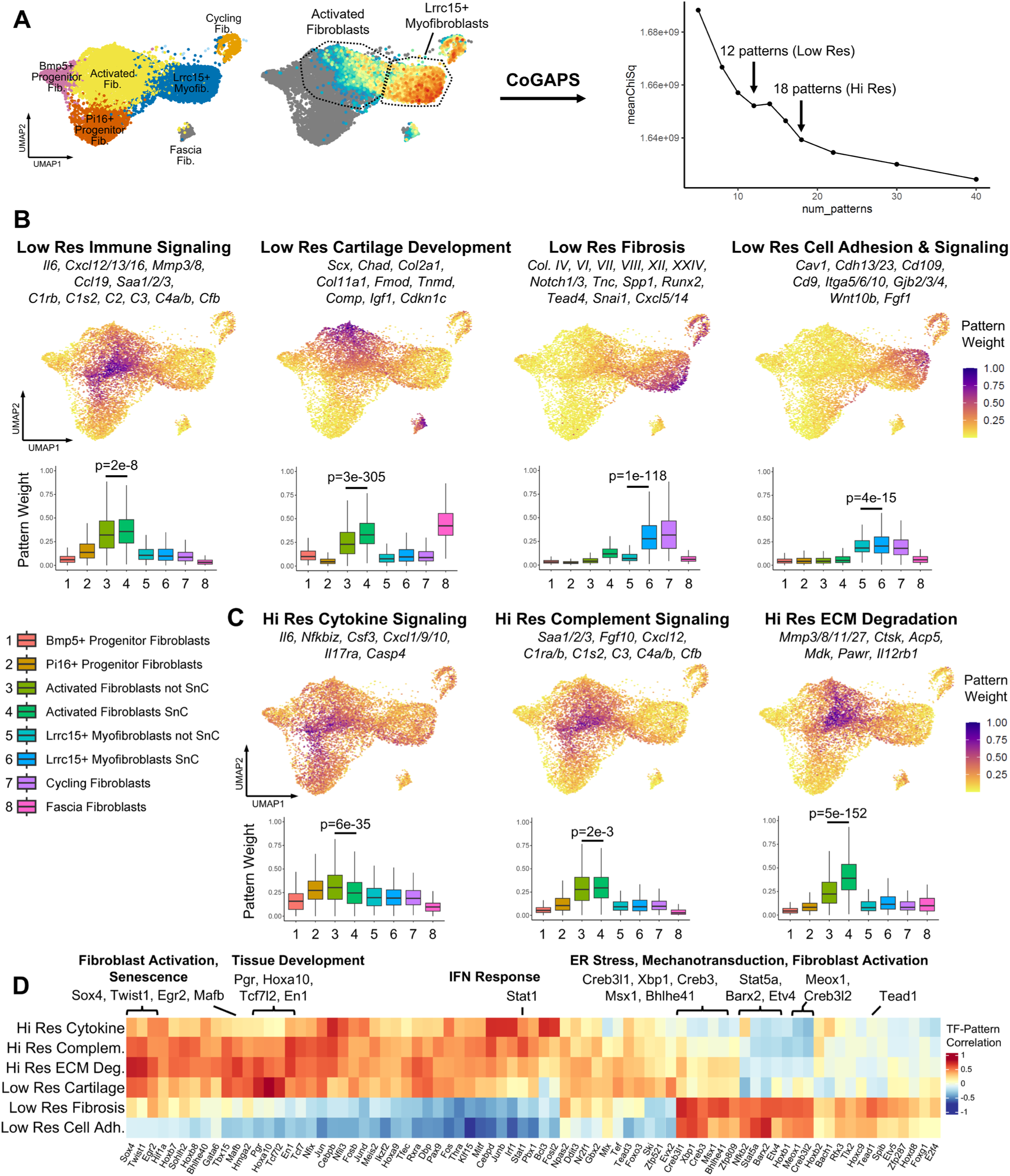
CoGAPS analysis reveals biological patterns associated with specific fibroblast SnC subtypes. **A)** Application of CoGAPS to all fibroblast subclusters (left UMAP). SenSig projection weight, originally displayed in Fig. 1A, is shown for reference (right UMAP). The optimal number of patterns was determined by plotting of the pattern number (dimension) and meanChiSq value for each CoGAPS run. The *k =* 12 (Low Res) and *k* = 18 (Hi Res) dimensions were selected for further analysis based on their drop in meanChiSq. **B)** Low Res CoGAPS Patterns representing different fibroblast SnC subpopulations. **C)** Hi Res CoGAPS Patterns that indicate different gene co-expression modules related to the Low Res CoGAPS “Immune Signaling” pattern. **B** and **C** show pattern weights in each cluster via UMAP and box plots. Select genes in each pattern are shown above the UMAP plots. A two-tailed t-test with Benjamini-Hochberg correction between the pattern weights of SnCs and non-SnCs within each cell type was used for the analysis in **B** and **C**. **D)** Spearman correlation coefficient between TF activity and CoGAPS patterns. TF activity was computed using pySCENIC^46^. Only the TFs with a correlation >0.3 in at least one of the patterns are shown in the plot. Correlations with a p-value <0.05 were set to 0.

Using the Low Resolution CoGAPS, we identified senotype gene co-expression patterns that reflect functionally diverse fibroblast states (**Fig. 3B**, **Ext. Data 5A, Supplemental Data 4**). There were four patterns that were upregulated in different SnC subpopulations (called senotype-specific patterns) and one pattern was expressed across all SnCs (**Fig. 3B**, **Ext. Data 5A**). The pattern spanning all fibroblast SnCs included co-expression of genes involving cell cycle arrest, such as *Cdkn2a* and *Trp53inp1*, and cellular stress—processes known to be upregulated in SnCs. This suggests that all fibroblast senotypes express common pathways related to cell cycle arrest and senescence induction. By contrast, the other four senotype-specific patterns were expressed in distinct fibroblast subpopulations and represented different biological pathways (**Fig. 3B**).

Myofibroblasts expressed two senotype patterns that exhibited transcriptional programs related to fibrotic signaling and cell adhesion. We annotated the first senotype-specific pattern as “Fibrosis” because it included co-expression of genes involving collagen and basement membrane ECM production, fibrotic signaling, Notch signaling, and epithelial mesenchymal transition, such as *Tnc*, *Spp1*, *Notch1/3*, and *Snai1* (**Fig. 3B**). We named the second senotype-specific pattern “Cell Adhesion & Signaling” due to its co-expression of genes for integrins, gap and tight junction proteins, intermediate filaments, and exosome membrane proteins, indicating a phenotype involving cell adhesion and signaling rather than ECM production (**Fig. 3B**). The cell weights for both patterns correlated with the activity of transcription factors (TFs) estimated independently with SCENIC^46^ such as *Cre3bl1*, *Xbp1*, *Creb3*, and *Creb3l2*, suggesting that these myofibroblast senotypes both undergo ER stress (**Fig. 3D**). However, the Fibrosis pattern activated unique TFs related to mechanoresponsive and EMT-associated TFs such as *Tead1*, *Etv4*, and *Etv5*, while the Cell Adhesion & Signaling pattern was more correlated with *Stat5a* and *Barx2*—TFs associated with cell adhesion and differentiation. These data suggest myofibroblasts express gene programs involving two key aspects of fibrosis development: one related to fibrotic ECM production and the other related to cell-ECM interactions and ECM organization.

Activated fibroblasts expressed the other two senotype patterns representing distinct cartilage development and immune signaling programs. One of these patterns was distinguished from the others based on its co-expression of cartilage- and tendon-associated genes, including *Col2a1*, *Col11a1*, *Fmod*, *Scx, Mgp*, and *Comp*, leading to annotation as the “Cartilage Development” senotype (**Fig. 3B**). In addition, this pattern co-expressed genes related to mechanotransduction and Wnt signaling, such as *Piezo2*, *Ccn2*, and *Lgr5*. Furthermore, this pattern correlated with the activity of TFs related to Wnt signaling and skeletal development, such as *Tcf7l2*, *Hoxa10*, and *Evx2* (**Fig. 3D**). These data suggest this senotype activates an alternative pro-fibrotic pathway distinct from myofibroblast senotypes that leads to the deposition of fibrocartilage ECM. By contrast, the other senotype pattern we annotated as “Immune Signaling,” co-expressed various cytokines, chemokines, complement factors, and ECM degradation enzymes known to be involved in inflammation and tissue remodeling (**Fig. 3B**). Notably, this was the only senotype pattern that was positively correlated with SenMayo^39^ (**Ext. Data 5D**), suggesting this pattern captures a pro-inflammatory and ECM remodeling phenotype that resembles the senotypes identified in other tissues and disease contexts.

We also found that the Immune Signaling pattern was expressed at the boundary of senescence, while the other senotype patterns were predominantly expressed by SnCs. Therefore, we hypothesized that the genes defining the Immune Signaling senotype could be refined using a higher dimension of CoGAPS. In evaluating the Chi-squared fit from CoGAPS used to evaluate dimensionality, we observed a second drop in the fit at *k* = 18 (**Fig. 3A**). Our previous studies have shown that different values of *k* resolve distinct biological features consistent with cellular hierarchies inferred with multi-dimensional factorizations^47–49^. Therefore, we decided to explore the co-expression signatures and associated cell weights defining the patterns learned from the CoGAPS factorization at the *k* = 18 dimension, which we refer to as “High Resolution CoGAPS” (**Fig. 3B-C**, **Ext. Data 5A-C, Supplemental Data 4**).

From the High Resolution CoGAPS, we found three different patterns that were highly expressed by the Immune Signaling senotype related to other fibroblasts (**Ext. Data 5B**), but only two of those patterns were upregulated by SnCs relative to non-SnCs. The pattern present in non-SnCs, which we named “Cytokine Signaling,” included several signaling molecules traditionally recognized as SASP factors —such as *Il6*, *Cxcl1*, *Cxcl9*, *Cxcl10,* and *Csf3*—as well as *Il17ra*, a receptor previously linked to senescence induction^50^. This suggests that these cytokine signaling factors are not specifically expressed by the fibroblast SnCs in our fibrosis model. Instead, we observed an alternative subset of pro-inflammatory SASP factors that were expressed in this senotype, as discovered in the other two patterns. One of those patterns, which we named “Complement Signaling,” included *Cxcl12*, *Saa3*, and several complement-associated genes—genes known to be overexpressed in inflammatory fibroblasts in other disease contexts^51,52^. The other pattern, which we annotated as “ECM Degradation,” co-expressed genes related to ECM degradation, including *Mmp3*, *Ctsk*, and *Acp5*. Both the Complement Signaling and ECM Degradation patterns were correlated with TFs linked to fibroblast activation and senescence, such as *Sox4*, *Twist1*, *Egr2*, and *Mafb* (**Fig. 3D**). These data suggest that the Immune Signaling senotype engages in complement signaling, immune cell recruitment, and ECM remodeling as a result of persistent fibroblast activation.

Taken together, through hierarchical factorization, we discovered four distinct fibroblast senotypes that result from a common senescence program involving cellular stress and cell cycle arrest. In our remaining analyses, we refer to these senotypes by names representing their associated Low Resolution CoGAPS patterns: Fibrosis, Cell Adhesion & Signaling, Cartilage Development, and Immune Signaling. In addition, we explore the High Resolution CoGAPS Immune Signaling sub-patterns in the following section to infer senotype trajectory.

### Fibroblast senotypes evolve across fibrosis development

Recent studies have shown that fibroblasts have the potential to differentiate along variable trajectories during wound healing and fibrosis development^51,53,54^. *In vitro* evidence also suggests that SnCs may follow distinct trajectories following induction^55^. However, it remains less understood whether fibroblast SnCs follow distinct trajectories *in vivo*. To explore the trajectory of fibroblast senotypes across fibrosis development, we applied projectR transfer learning of the broad SenSig for SnC identification and the refined senotypes defined from our low and high resolution CoGAPS patterns to our previously published scRNA-seq data of sorted CD29+CD45-CD31-from a VML injury^25^. This dataset includes cells isolated from uninjured muscle and VML injuries treated with saline, decellularized ECM, or PCL at weeks 1 and 6 following treatment (**Ext. Data 6A**), enabling us to determine the kinetics of senotypes involved in wound healing and fibrosis development. We assigned cluster names congruent with the annotations in CD45-enriched scRNA-seq object to facilitate comparisons between the datasets (**Ext. Data 6B**). Using transfer learning with SenSig, we found that two fibroblast clusters, activated fibroblasts/myofibroblasts and cartilage-like fibroblasts, exhibited high SenSig projection weights (**Ext. Data 6C**), as documented previously^25^.

Among these two clusters, we found that the fibroblast senotypes emerged at different stages of wound healing and fibrosis development (**Ext. Data 6D-I**). The projection weights of both myofibroblast patterns, Fibrosis and Cell Adhesion & Signaling, were elevated in all groups at Week 1, but they declined by Week 6 in all groups except the PCL (fibrotic) condition (**Ext. Data 6D-E**). By contrast, Cartilage Development and Complement Signaling projection weight peaked at Week 6 and was highest in the PCL condition (**Ext. Data 6F, H**). Similarly, ECM Degradation projection weight was elevated at Week 6 compared to Week 1 in the PCL- and decellularized ECM-treated groups, but it was not present in the saline-treated group (**Ext. Data 6I**). Furthermore, the highest ECM Degradation projection weight was observed in the decellularized ECM, a treatment that results in muscle regeneration rather than fibrosis^56^.

These findings demonstrate that fibroblasts evolve during the progression of fibrosis. Myofibroblast senotypes emerge during acute wound healing but persist through conditions of chronic inflammation and fibrosis (PCL). Comparatively, Cartilage Development, Complement Signaling, and ECM Degradation fibroblast senotypes emerge only during the chronic inflammation and fibrosis stages (PCL). The ECM Degradation senotype is also present during regenerative processes (decellularized ECM), suggesting this senotype may play a broader role in tissue remodeling outside the fibrotic context.

### Senotypes occupy distinct cellular and ECM niches

Differences in gene expression profiles of fibroblast and perivascular senotypes suggest they may occupy different niches within the fibrotic tissue. To map senotype niches, we performed VisiumHD on a fibrotic tissue section generated using the same injury and biomaterial treatment as scRNA-seq (**Fig. 1A**), enabling an integrated analysis between the two datasets. To identify the different cell subpopulations within the VisiumHD dataset, we performed clustering at the 8 μm bin resolution using the Seurat v5 pipeline^27^. We used this approach to enable identification of cell types that were absent from our scRNA-seq dataset due to technical artifacts from tissue digestion.

This analysis revealed clusters resembling those from the scRNA-seq dataset, including fibroblasts, vascular cells (ECs and perivascular cells), lymphatic vessels, myeloid cells, fascia fibroblasts, and skeletal muscle cells (**Ext. Data 7A**, **Supplemental Data 5**). We also identified clusters containing multiple cell types, suggesting select cell subpopulations coexist in the same spatial niche. One of these clusters contained lymphocytes (B cells and T cells), antigen-presenting myeloid cells, and activated fibroblasts, while the other included *Pi16*+ progenitor fibroblasts and *Mrc1*-high myeloid cells. We mapped these clusters onto the accompanying H&E image to determine the spatial distribution of cell subpopulations relative to the implant and surrounding tissues. This process revealed that the *Pi16*+ fibroblast and *Mrc1*-high myeloid cell cluster was located along the exterior, adipose-facing side of the implant, while the fascia fibroblasts were in the fascia and at the muscle-implant interface (**Ext. Fig. 7A**). The other major clusters were interspersed between the implant particles, suggesting these cell types are the predominant subpopulations contributing to inflammation and fibrosis around the implant.

Next, we mapped senotypes by conducting projectR transfer learning with SenSig and the fibroblast CoGAPS patterns from the *k* = 12 dimension (**Fig. 4A-C**, **Ext. Data 7B**). Through this analysis, we identified spatial niches linked to different senotypes. Myofibroblast senotypes were found alongside myeloid cells near the implant particle surface, a region we named the “SnC Fibrotic Signaling” niche (**Fig. 4A**). This niche contained high expression of fibrotic ECM and signaling genes, including *Col6a1*, *Tnc*, and *Spp1* (**Fig. 4A**). Immune Signaling fibroblast senotypes resided further away from the implant surface—colocalizing with clusters containing fibroblasts, antigen-presenting myeloid cells, and lymphocytes, while expressing higher levels of *Mmp3*, *Saa3*, and *Cxcl12* relative to other regions (**Fig. 4B**). We named this region the “SnC Immune Signaling & Tissue Remodeling Niche.” Cartilage Development fibroblast senotypes were adjacent to both the immune and fibrotic signaling niches and did not colocalize with other immune cells. This region had higher levels of pro-fibrotic genes such as *Col1a1* and cartilage-associated genes such as *Mgp* and *Cilp* (**Fig. 4C**). Therefore, we annotated this region as the “SnC Cartilage Development” niche. Lastly, vascular SnCs appeared in both the SnC Fibrotic Signaling and Immune Signaling & Tissue Remodeling niches in proximity to fibroblast SnCs (**Fig. 4A, B**).

**Figure 4.**
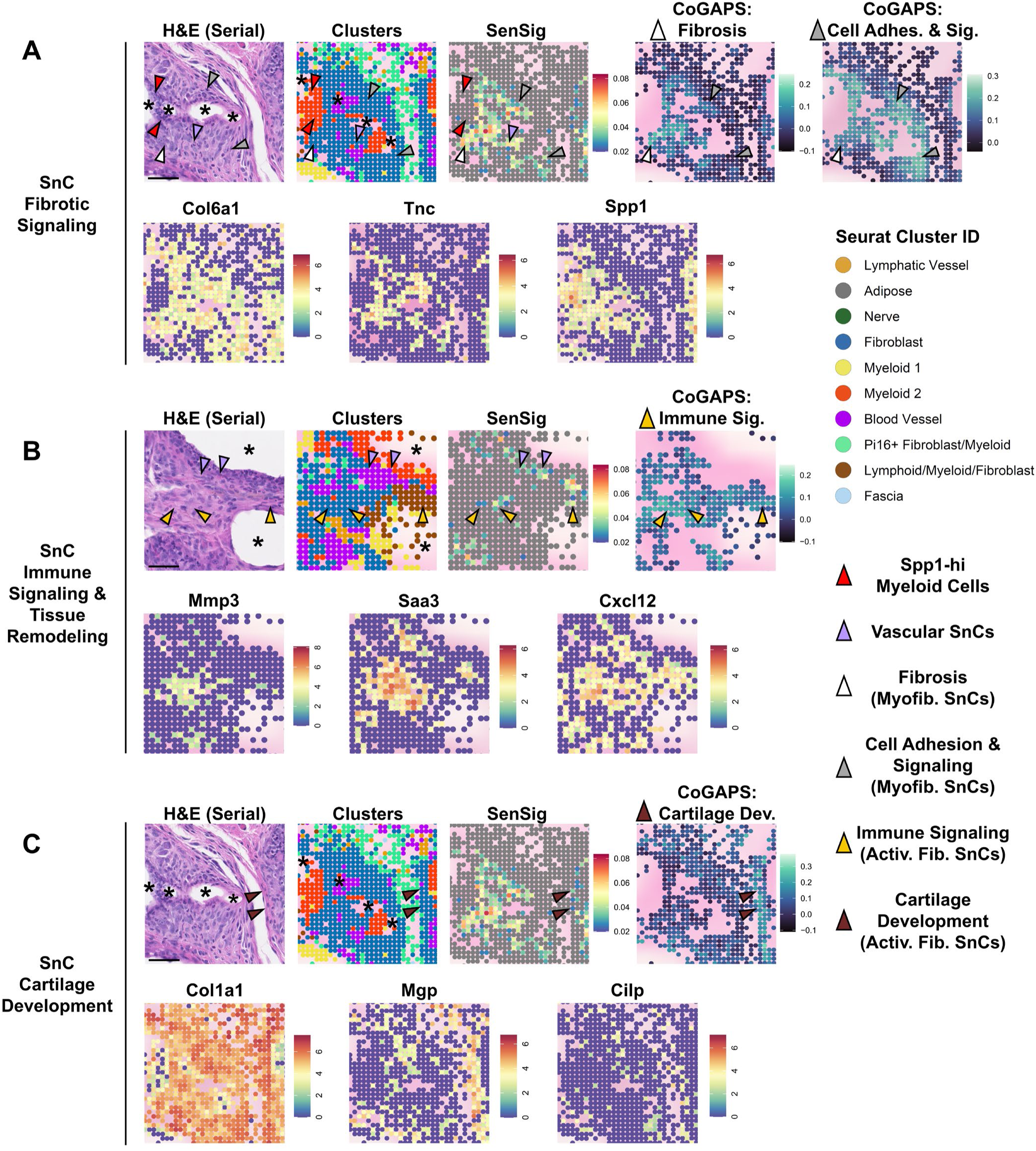
VisiumHD analysis reveals spatial niches associated with specific senotypes. **A-C)** Detection of Fibrotic Signaling, Immune Signaling & Tissue Remodeling, and Cartilage Development senotype niches. (Top Row) H&E images (serial section, scale bar: 50 µm), VisiumHD clusters, SenSig projection (p < 0.01), and fibroblast CoGAPS pattern projections in the fibrotic implants. CoGAPS pattern projections are only shown in clusters containing fibroblasts. Arrows highlight regions where each senotype is present in the tissue. (Bottom Row) Spatial feature plot showing normalized expression of senotype-specific genes in each niche. Asterisks (*) indicate the location of the PCL particles in the H&E and VisiumHD sections.

To determine whether these senescent niches are coupled with different tissue microarchitectures, we performed high-resolution imaging of the serially sectioned H&E-stained tissue and identified the regions congruent to the VisiumHD section. The SnC Fibrotic Signaling niche exhibited high cell density, and its ECM had a less defined fibrillar architecture than surrounding regions (**Fig. 4A**). By contrast, the SnC Immune Signaling & Tissue Remodeling niche contained loosely organized ECM fibrils—a finding consistent with the high expression of ECM degradation genes observed in the immune signaling fibroblast senotypes (**Fig. 4B**). Lastly, the SnC Cartilage Development niche contained ECM fibers resembling the structure of muscle fascia and tendon (**Fig. 4C**).

To probe the cell-cell interactions driving niche development, we implemented a second approach to label each 8 μm bin using RCTD^57^, a package that performs bin-level cell type decomposition using annotations obtained through a reference scRNA-seq object. We used this approach to better delineate bins containing a mixture of cell types, such as fibroblasts or myeloid cells, that could not be obtained through unsupervised clustering. By applying RCTD using cell type labels obtained from scRNA-seq, we observed a high rate of doublet bins containing fibroblasts, myeloid cells, and vascular cells (**Ext. Data 8A-C**). These findings suggest that the higher rate of doublets is not merely a technical artifact; rather, it reflects that these cells may co-occur and interact with each other in distinct niches.

Using a refined clustering analysis, we found that fibroblast and perivascular senotypes spatially co-occur with different immune, stromal, and vascular cell populations. We clustered the RCTD-labeled fibroblasts and myeloid cells to obtain a higher resolution of cell subpopulations present in the fibrotic implant. Through this higher resolution clustering, we identified four relevant clusters with a comparatively elevated SenSig projection weight and *Cdkn2a* expression: Three containing fibroblasts and one containing vascular cells. (**Ext. Data 8D-F, Supplemental Data 5**). One fibroblast cluster expressed marker genes for activated fibroblasts, B cells, and antigen-presenting myeloid cells, indicating that it represents the SnC Immune Signaling & Tissue Remodeling niche (**Ext. Data 8D**). The other two fibroblast clusters had a high projection weight of the myofibroblast CoGAPS patterns, suggesting that they represent the SnC Fibrotic Signaling niche (**Ext. Data 8G**). These clusters expressed marker genes for myofibroblasts, activated fibroblasts, vascular cells, macrophages, and neutrophils, suggesting these cell types spatially co-occur in this niche (**Fig. 4A**, **Ext. Data 8D-E**). Additionally, select bins within these clusters had an elevated Immune Signaling pattern weight (**Ext. Data 8G**), consistent with our observation that the SnC Immune Signaling & Tissue Remodeling niche occurs in regions separate from, but sometimes adjacent to, the SnC Fibrotic Signaling niche (**Fig. 4B**). All three fibroblast clusters contained bins with high expression of the Cartilage Development pattern (**Ext. Data 8G**), supporting our finding these Cartilage Development fibroblast SnCs are adjacent to the SnC Fibrotic Signaling and Immune Signaling & Tissue Remodeling niches (**Fig. 4C, Ext. Data 7B**). Finally, the vascular cell cluster co-occurred to a limited extent with clusters that are part of the SnC Fibrotic Signaling and Immune Signaling & Tissue Remodeling niches (**Ext. Data 8H**). Perivascular SnC genes were observed in both the vascular cell cluster and the SnC Fibrotic Signaling clusters. (**Ext. Data 8D**). These findings suggest perivascular SnCs are confined to the interfaces of the SnC Fibrotic Signaling and Immune Signaling & Tissue Remodeling niches.

To elucidate the interactions driving the development of the senotype niches, we evaluated cell-cell signaling using Squidpy^58^ co-occurrence analysis and SASP factor annotation of differentially expressed genes in each senotype-associated cluster. The two clusters associated with the SnC Fibrotic Signaling niche spatially co-occurred with each other and with a cluster containing neutrophils and inflammatory macrophages (**Ext. Data 8H**). These clusters expressed higher levels of fibrotic and immunosuppressive SASP factors *Spp1*, *Cxcl5*, and *Tgfb1*—all of which were found adjacent to receptors expressed by neutrophils, macrophages, or fibroblasts (**Ext. Data 9A-C**). Perivascular senotype SASP factors associated with vascular and fibrotic remodeling, such as *Pdgfa*, *Serpine2*, *Igfbp7*, *Col4a1*, and *Col18a1*, were also found in these clusters and the vascular cell cluster (**Ext. Data 9B**). In particular, *Pdgfa*+ regions co-localized with *Pdgfra*, a receptor that leads to fibroblast activation when bound to PDGF-AA—the protein that corresponds with the *Pdgfa* gene (**Ext. Data 9C**). These data suggest that the myofibroblast and perivascular SnCs engage in signaling with myeloid cells, fibroblasts, and vascular cells within the SnC Fibrotic Signaling niche, leading to immunosuppression, vascular remodeling, and fibrosis development.

By contrast, signaling interactions within the SnC Immune Signaling & Tissue Remodeling niche were associated with immune cell recruitment and activation. The cluster associated with this niche spatially co-occurred with other lymphocyte-containing clusters, rather than the clusters associated with fibrotic signaling (**Ext. Data 8H**). In addition, this cluster expressed higher levels of immune signaling fibroblast SASP factors *Cxcl12* and *Saa3* (**Ext. Data 9B**), which were co-localized with *Cxcr4* and *Tlr2*—receptors expressed by the innate and adaptive immune cells within this niche (**Ext. Data 9C**). *Cxcl12*+ bins also co-localized with another receptor, *Ackr3* (**Ext. Data 9C**). However, this receptor was expressed by fibroblasts and lymphatic ECs located in different regions compared to *Cxcr4*+ immune cells (**Ext. Data 9C**). CXCL12-CXCR4 binding is known to promote immune cell recruitment, while ACKR3 acts as a scavenger receptor that limits CXCL12 signaling. Expression by these receptors in different cell types and spatial regions suggests the effects of CXCL12 signaling depend on the surrounding microenvironment.

Taken together, our findings illustrate that the senotypes defined by scRNA-seq are regionally confined to distinct functional units within the fibrotic tissue. Myofibroblast senotypes engage in fibrotic and immunosuppressive signaling with macrophages and neutrophils in cellular-dense regions (SnC Fibrotic Signaling niche), while the Immune Signaling fibroblast senotype promotes immune cell activation and matrix degradation in zones of loosely-organized ECM (SnC Immune Signaling & Tissue Remodeling niche). The Cartilage Development senotype is regionally confined to immune-excluded, ECM-dense regions (SnC Cartilage Development niche) that surround the other two niches. Perivascular SnCs span the SnC Fibrotic Signaling and Immune Signaling & Tissue Remodeling niches, suggesting that these cells engage in processes that bridge immune signaling, vascular remodeling, and fibrosis development.

### Perivascular senotypes express inflammatory response and ECM remodeling pathways

Through spatial transcriptomics and multiplex IF, we found that senescent perivascular cells localize to SnC-dense regions spanning both the Fibrotic Signaling and Immune Signaling & Tissue Remodeling niches (**Fig. 4A-B**, **Fig. 5A**). Although less abundant than fibroblast senotypes, their localization at the interface of these niches suggests that perivascular SnCs may participate in coordinating inflammatory responses, vascular remodeling, and fibrosis development. To define the molecular programs associated with these cells, we performed gene set enrichment analysis comparing senescent and non-senescent perivascular cells within the scRNA-seq dataset (**Fig. 1A**). Perivascular SnCs upregulated pathways related to epithelial mesenchymal transition and ECM organization (**Fig. 5B**), including genes for various fibrillar collagens (*Col3a1, Col5a1*), basement membrane ECM (*Col4a1/2, Col18a1, Lama2/4*), and ECM remodeling enzymes (*Adamts2/9, Adam12, Mmp14*) (**Ext. Data 10A**). These SnCs also upregulated cytokine and stress responses pathways, including genes associated with NF-κB activation (*Nfkb1, Nfkb2, Tnfaip2, Tnfaip3*), interferon response (*Ifngr1/2, Ifitm1/2, Stat1*), cytokine signal transduction (*Il1r1, Il6st*), and cellular stress (*Ddit4, Hif1a*) (**Fig. 5B, Ext. Data 10B**).

**Figure 5.**
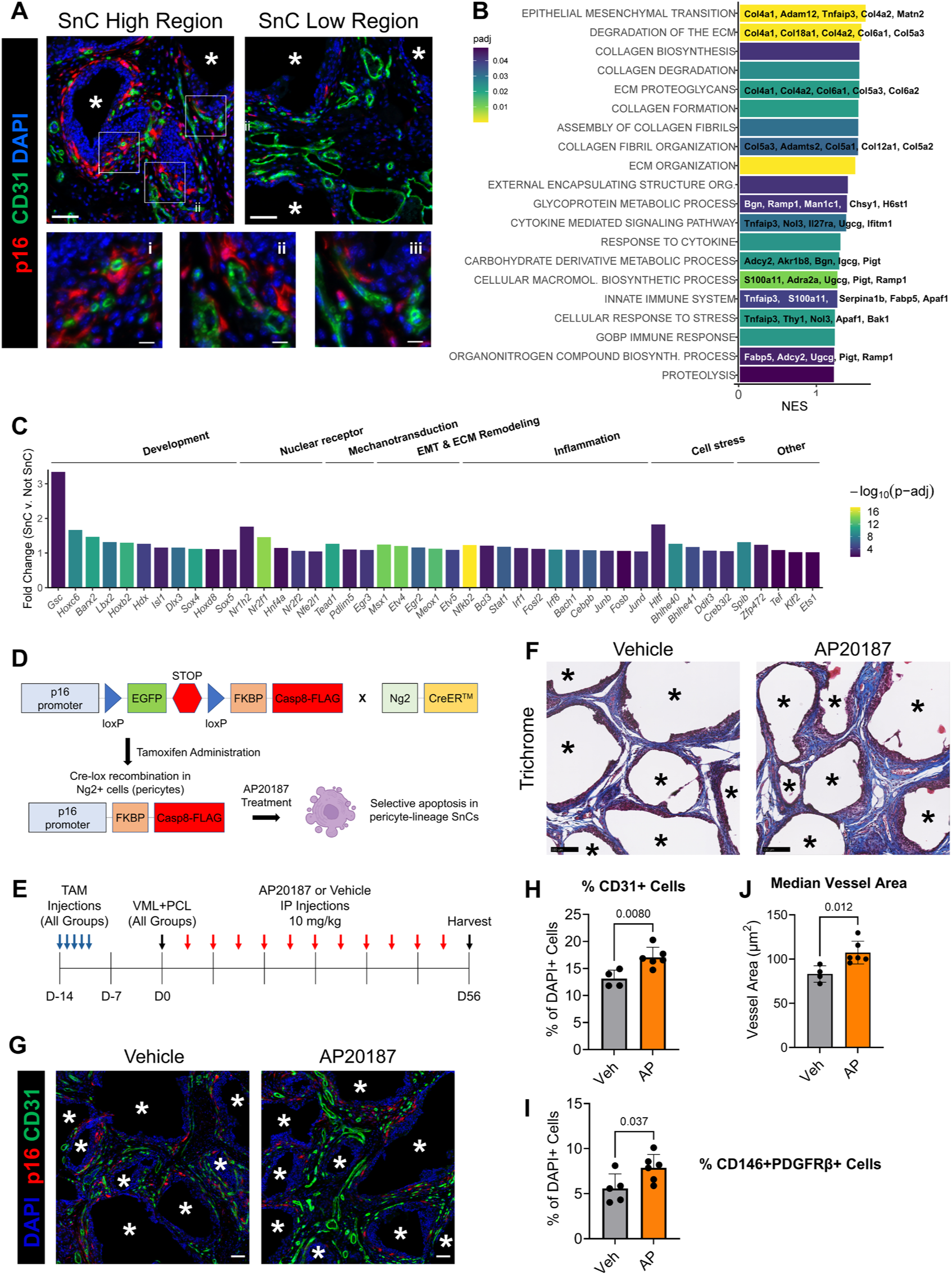
Perivascular SnCs modulate vascularization and fibrosis in response to VML and synthetic biomaterial implantation. **A)** IF image showing p16+ cells and CD31+ ECs in regions high and low in SnCs. Perivascular p16+ cells are observed nearby capillaries SnC-high regions (i – iii). Scale bar: 50 µm (top), 10 µm (bottom) **B)** Significantly enriched GSEA pathways from the HALLMARK, REACTOME, and GOBP databases (p_adj_<0.05). The top leading edge genes are shown for select pathways. C) Differentially expressed regulons (p_adj_<0.05) in perivascular SnCs versus Not SnCs. Regulon activity (AUC) was computed using pySCENIC on the entire scRNA-seq dataset shown in Fig. 1A. Fold change values represent (Median AUC SnC)/(Median AUC Not SnC). TFs are arranged according to their associated biological categories listed at the top of the plot. **D)** Transgenic murine model to selectively eliminate pericyte-lineage SnCs. **E)** Treatment regimen for pericyte-lineage SnC deletion. **F-G)** Trichrome (**F**) and immunofluorescence staining (**G**) showing higher amounts of collagen and CD31+ blood vessels with pericyte-lineage SnC elimination. Scale bar: 100 µm (**F**), 50 μm (**G**) **H-**Percentage of CD31+ cells (ECs, **H**), percentage of CD146+PDGFRβ+ cells (pericytes, **I**), and median area of CD31+ vessels (**J**). Vessels with an area <20 μm^2^ were excluded from analysis. Data are mean ± standard deviation. n=4-6 biological replicates, Two-tailed t-test. Asterisks (*) indicate regions with PCL particles in **A**, **F**, and **G**.

Using SCENIC, we identified activated TFs (regulons) in perivascular SnCs that connect inflammatory responses to ECM remodeling pathways. Regulons upregulated in perivascular SnCs compared to non-SnCs included those involving response to inflammatory cytokines (*Stat1, Nfkb2, Bcl3, Irf1/8, Cebpb, Fosb, Fosl2, Junb/d*), response to cellular stress (*Bach1, Bhlhe40/41, Ddit3, Creb3l2, Hltf*), vascular regulation (*Tead1, Pdlim5, Nr2f1/2*), and EMT and ECM remodeling (*Meox1, Msx1, Gsc, Sox4/5, Etv4/5, Barx2, Egr2*) (**Fig. 5C**). Notably, some of the regulons related to cytokine response and cellular stress, such as *Cebpb*, *Creb3l2*, *Junb*, *Irf1/8*, and *Nfkb2*, included downstream genes associated with EMT and vascular and ECM remodeling (**Ext. Data 10C**). These data suggest perivascular SnCs respond to inflammatory cues and, in turn, upregulate SASP related to vascular and ECM remodeling.

### Pericyte-lineage SnCs modulate vascular and fibrosis development

Due to perivascular SnCs’ involvement in diverse processes related to vascular remodeling and fibrosis development, we devised an approach to selectively eliminate perivascular SnCs in order to discern their functional roles. We combined the Ng2-CreER™ and p16-LOX-ATTAC strains to enable selective elimination of pericyte-lineage cells (**Fig. 5D**). The Ng2-CreER™ model enables selective expression of Cre recombinase in pericytes and has been successfully applied to selectively track and target pericyte-lineage cells in cardiac muscle fibrosis models^59,60^. The p16-LOX-ATTAC model enables Cre-inducible expression of the FKBP-Caspase8 “suicide” gene in p16-expressing cells. Administration of AP20187 dimerizes the FKBP-Caspase8 protein to activate apoptosis, enabling targeted elimination of specific types of p16-expressing cells. This model was recently developed and applied to eliminate senescent osteocytes^61^ and ECs^62^ in murine aging and obesity models, respectively, illustrating its utility to study different SnC subpopulations in multiple disease contexts.

To eliminate pericyte-lineage SnCs, we first administered tamoxifen two weeks prior to VML and biomaterial implantation to enable Cre recombination of the p16-LOX-ATTAC allele in steady state pericytes (**Fig. 5E**). We then administered AP20187 twice weekly to deplete pericyte-lineage p16+ cells throughout the fibrosis progression (**Fig. 5E**).

Twice weekly depletion of pericyte-lineage SnCs affected vascular maturation and fibrosis progression. In the group with pericyte-lineage SnC depletion, there was qualitative increases in collagen staining and quantitative increases in CD31+ cell (EC) and CD146+PDGFRβ+ cell (pericyte) percentage around implant particles, but the percentage of p16+ cells, p16+CD31+, and p16+ CD146+PDGFRβ+ cells did not significantly change (**Fig. 5F-I, Ext. Data 10D-F, Ext. Data 11A-B**). Furthermore, the median vessel area and the proportion of larger vessels increased in the pericyte-lineage SnC-depleted group (**Fig. 5J**, **Ext. Data 10G**). These findings suggest that pericyte-lineage SnCs prevent the development of mature blood vessels and attenuate fibrosis progression rather than contributing to fibrotic matrix deposition.

### Fibroblast and perivascular senotypes are present in other murine and human fibrotic conditions

To query fibroblast and perivascular cell senotypes across other datasets, we developed SENgEAR, a specialized instance of the gEAR (Gene Expression Analysis Resource) platform^19^. SENgEAR enables users to interactively explore and analyze the datasets described here alongside a broader collection of curated public transcriptomic datasets, including both single-cell and spatial transcriptomics studies relevant to cellular senescence. Through an intuitive web interface, users can query gene expression, compare patterns across cell types and conditions, and evaluate senescence-associated gene signatures across studies. By integrating the senotype signatures discovered through our analyses with external resources in a unified analytical environment, SENgEAR facilitates reproducibility, cross-study comparison, and custom exploration in the research community.

We used a collation of public datasets and SENgEAR to investigate whether the senotypes discovered in our fibrosis model are present in other fibrotic conditions. First, we applied the SENgEAR pipeline to a scRNA-seq dataset of murine pulmonary fibrosis. This dataset was derived from sorted mesenchymal cells in a murine bleomycin-induced lung injury model^51^. In previous work, this dataset was used to trace the pro-fibrotic lineage of alveolar fibroblasts, which differentiate into fibrotic fibroblasts through an intermediate inflammatory fibroblast state^51^. However, it remains unclear whether these fibroblast subpopulations become senescent, and if so, which senotypes they express.

By performing transfer learning of SenSig and fibroblast senotype CoGAPS patterns through SENgEAR, we found that several fibroblast senotypes are present in bleomycin-induced pulmonary fibrosis. Transfer learning of SenSig identified cells with a high senescence score in the fibrotic cluster—the cluster containing the predominant cell population contributing to fibrosis^51^ (**Fig. 6A, Ext. Data 12A**). We observed a higher projection weight of the Cell Cycle Arrest pattern and higher *Cdkn2a* expression in this cluster, supporting the SenSig-based senescence prediction (**Ext. Data 12B, C**). Transfer learning of the CoGAPS patterns revealed that the Fibrosis and Cartilage Development pattern weights were highest in the fibrotic cluster, but their weights varied between different cell subpopulations (**Fig. 6A**). Cells with the highest Fibrosis pattern weight were present in both the inflammatory and fibrotic fibroblasts. These cells expressed pattern marker genes including *Spp1*, *Serpine2*, *Tnc*, and *Cxcl14* (**Fig. 6A**, **Ext. Data 12E**). By contrast, cells with the highest Cartilage Development pattern weight were only present in the fibrotic cluster and expressed fibrocartilage genes such as *Fmod*, *Col11a1*, *Scx*, and *Igf1* (**Fig. 6A**, **Ext. Data 12F**). We also observed a high pattern weight of the Cell Adhesion and Signaling pattern in a smaller subset of the fibrotic fibroblasts (**Fig. 6A**). This cell subpopulation expressed cell adhesion and junction genes such as *Itga5*, *Gjb3/4/5*, and *Cav1* (**Ext. Data 12G**).

**Figure 6.**
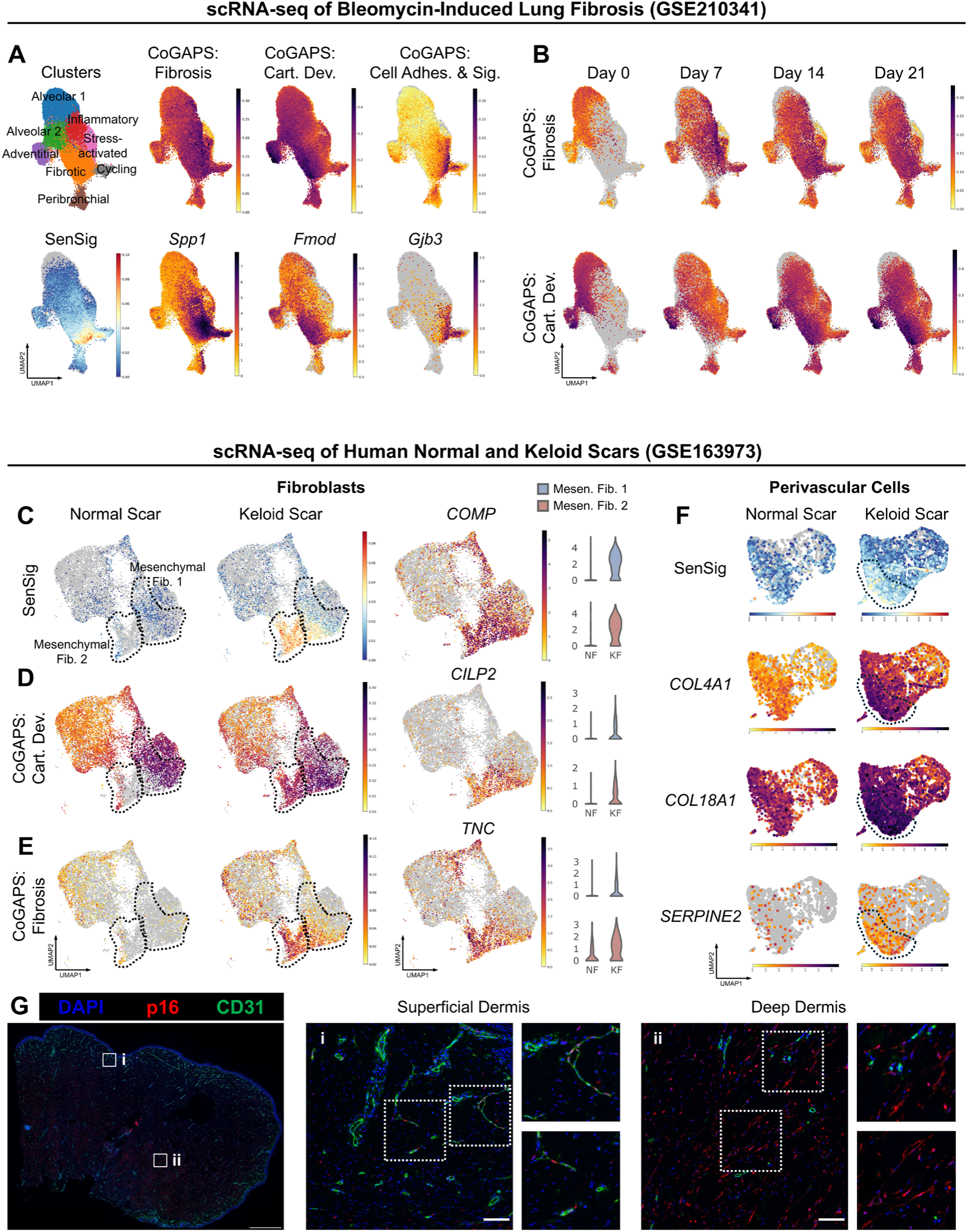
Fibroblast and perivascular senotypes are present in other fibrotic tissues. **A-B)** Senotype identification in bleomycin-induced lung fibrosis. Transfer learning of SenSig and Low Resolution CoGAPS patterns was performed on a dataset of fibroblasts isolated from a bleomycin-induced lung injury at Days 0 (no injury), 7, 14, and 21 (GSE210341). **A** shows UMAP projection of fibroblast clusters, SenSig projection weight, and CoGAPS projection weight. Normalized expression of select CoGAPS pattern marker genes are shown below their respective pattern. **B** shows projection weight of Fibrosis and Cartilage Development patterns at each timepoint. **C-F)** Senotype identification in human keloid scars. Transfer learning of SenSig (**C, F**) and CoGAPS (**D-E**) patterns was performed on a dataset of cells isolated from patient-matched normal scars (NF) and keloid scars (KF) (GSE163973, n=3 patients). **C** shows projection weight of SenSig and normalized expression of *COMP*, a SASP factor detected across all SenSig-high cells in keloid scars. The two mesenchymal clusters are outlined in black. **D** and **E** show the projection weight of CoGAPS patterns detected in keloid SnCs: Cartilage Development (**D**) and Fibrosis (**E**). Normalized expression of pattern-specific SASP factors that are expressed in keloid scars is shown on the right. **F** shows SenSig projection weight and normalized gene expression of SASP factors in perivascular cells. SenSig-high cells are outlined in black. **G)** Multiplex IF images of human keloid tissue section showing p16+ cells with perivascular and fibroblastic morphologies. The brightness of the left-most image was increased by 40% using Microsoft Powerpoint to enable visualization of the full tissue. Scale bar: 1000 µm (left), 50 µm (right; i and ii)

Next, by evaluating the CoGAPS pattern weights at different timepoints, we determined that these senotypes in bleomycin-induced lung fibrosis exhibit a similar trajectory to those in our muscle fibrosis model. To map senotype trajectories, we evaluated the fibroblast CoGAPS pattern weights at the Day 0 timepoint (no injury), as well as Days 7, 14, and 21 following a bleomycin-injured injury. Day 7 represents the acute phase of injury, characterized by the presence of pro-inflammatory fibroblasts, while Days 14 and 21 represent the fibrotic phases. From this analysis, we found that the projection weight of each CoGAPS pattern varied with time. The Fibrosis pattern peaked at the Day 7 timepoint in inflammatory and fibrotic fibroblasts and remained elevated at Days 14 and 21 in the fibrotic fibroblasts (**Fig. 6B**). In addition, the Cell Adhesion and Signaling pattern was elevated at all timepoints in the fibrotic cluster (**Ext. Data 12D**). By contrast, the Cartilage Development pattern did not increase in the fibrotic cluster until Day 14 (**Fig. 6B**), the timepoint representing the onset of fibrosis. These findings mirror those from our muscle fibrosis model (**Ext. Data 6**), in which myofibroblast senotypes emerged during the acute injury response and remained present during fibrosis, while Cartilage Development fibroblast senotypes developed only in the fibrotic stages.

We next applied the SENgEAR pipeline to a dataset of normal and keloid scars derived from human skin biopsies^63^, and observed the presence of fibroblast and perivascular SnCs consistent with the findings in our mouse models. Keloids are raised, overgrown scars in the skin that are a result of dysregulated wound healing and excess collagen deposition. While studies have documented the presence of senescence biomarkers p16, p21, and SA-β-Gal in keloid tissues^64–66^, it is unclear which senotypes are present in keloids and how they contribute to aberrant scarring. To identify senotypes in keloid scars, we conducted transfer learning of the ortholog-mapped SenSig and CoGAPS fibroblast signatures onto the scRNA-seq dataset. Cells with a high SenSig score were found in fibroblasts and smooth muscle cells (i.e. perivascular cells) (**Ext. Data 13A-B**). Expression of *CDKN2A* and the CoGAPS Cell Cycle Arrest pattern was also higher in these populations (**Ext. Data 13C-D**), supporting the SenSig-based prediction of senescence.

Next, by evaluating the CoGAPS senotype pattern projections in the fibroblast subclusters of the human dataset, we observed myofibroblast and cartilage senotypes that are similar to our mouse models. Those subclusters included secretory papillary, secretory reticular, pro-inflammatory, and mesenchymal fibroblasts. SenSig projection weight and *CDKN2A* expression were highest in the mesenchymal fibroblasts (**Fig. 6C, Ext. Data 13E-F**), the subpopulations enriched in keloid versus normal scar. The most differentiated subcluster of mesenchymal fibroblasts had a high Fibrosis pattern weight, expressing higher levels of marker genes such as *COL8A1*, *TNC*, *SERPINE1*, and *SERPINE2* (**Fig. 6E, Ext. Data 13J**). These cell subpopulations also expressed higher levels of *ACTA2* and *TAGLN*, exhibiting a myofibroblast phenotype (**Ext. Data 13G-H**). By contrast, the senescent subpopulation from the other mesenchymal fibroblast cluster expressed the highest Cartilage Development pattern weight, upregulating genes such as *CILP2*, *IGF1*, and *ELN*. In addition, they did not express *ACTA2* or *TAGLN*, suggesting these SnCs are different from the myofibroblast SnCs (**Fig. 6D, Ext. Data 13K**). However, both of these fibroblast SnC subpopulations expressed some common fibrotic SASP factors, including *COMP*, *COL5A1*, *COL6A1*, *COL11A1*, and *MDK*, at a higher level compared to normal scars (**Fig. 6C, Ext. Data 13I, J**).

In addition to fibroblasts, the keloid contained a perivascular senotype that expressed SASP factors similar to those observed in our mouse model (**Fig. 6F**). To better elucidate the phenotype of these cells, we plotted SASP factors from our murine fibrosis model (**Fig. 2C**) in the human perivascular cell cluster. We observed elevated expression of vascular basement membrane and ECM remodeling factors in SenSig-high perivascular cells, including *COL4A1*, *COL18A1*, and *SERPINE2*—suggesting that these cells resemble the ECM remodeling phenotype observed in our murine fibrosis model (**Fig. 6F**).

By conducting immunostaining of p16 with CD31 to validate our findings, we confirmed that fibroblasts and perivascular SnCs were present in the keloid. Our analysis revealed that cells positive for p16 were present in the superficial and deep (fibrotic) dermis of the keloid, while no p16+ cells were observed in the epidermis (**Fig 6G**). However, the distribution and types of observed p16+ cells were different between the layers. The superficial region contained sparse p16+ cells that were predominantly perivascular. By contrast, p16+ cell density was significantly higher in the deep dermis of the keloid and consisted of both p16+ fibroblasts and perivascular cells—the former of which was more abundant. These data corroborate the scRNA-seq predictions that fibroblast and perivascular cells are senescent and suggest that each senotype emerges in different layers of the keloid scar.

## Discussion

In this study, we developed a multimodal pipeline to identify SnCs, profile their phenotypes, and map their spatial niches within a senescence-enriched murine fibrosis model. Using scRNA-seq, we found that several types of fibroblasts and perivascular cells were senescent in the fibrotic tissue. We validated our predictions using protein-based multiplex IF staining of senescence-specific and cell type-specific markers. This process is critical to confirming transcriptomics-based predictions because SenSig and other published senescence gene sets can capture other cell types that are not senescent, such as fibrocartilage or inflammatory cells. We then discovered a hierarchy of senotypes and their transcriptomics signatures using CoGAPS and differential expression analysis, which enabled us to identify the SASP factors and biological pathways upregulated in each senotype. Applying these signatures to a paired VisiumHD dataset, we mapped each senotype to distinct spatial niches and cell-cell communication networks. By applying this approach, we discovered nuanced senotype transcriptional programs and niches that are linked to different microenvironments within inflamed, fibrotic tissue. We disseminate these senotype signatures and datasets through a new user-friendly web-based data infrastructure, SENgEAR, that allows for ready querying of these senotype signatures across conditions and facilitates subsequent exploration of these senotypes across species, tissues, and biomedical conditions.

Our study identified SASP factors that were unique to specific subsets of fibroblast and perivascular SnCs, providing a high-resolution atlas of SASP factors that builds upon previous literature. Although SASP factors are known to be heterogeneous and cell type-dependent, they have traditionally been recognized as a combination of proinflammatory cytokines, chemokines, proteases, and growth factors. We found that some previously reported SASP factors^39,67,68^ were upregulated in at least one of the senotypes unveiled through our analysis, including *Cxcl5*, *Cxcl12*, *Ccl8*, *Serpine1*, *Serpine2*, *Igf1*, and *Mmp3*. However, the SnCs in our model predominantly upregulated SASP factors related to ECM production and tissue remodeling—and only a subset of fibroblast SnCs upregulated pro-inflammatory factors. These findings were supported by our analysis of murine pulmonary fibrosis and human keloid datasets, where most fibroblast and perivascular SASP factors were associated with ECM production and remodeling. Furthermore, some of the most commonly reported SASP factors^69^, including *Il6*, *Il8*, *Ccl2*, and *Gdf15*, were not upregulated in any SnCs observed in our fibrosis model or publicly analyzed datasets. Our findings resemble recent studies showing that senescent fibroblasts in fibrotic environments express an ECM-producing rather than pro-inflammatory phenotype^8,70,71^.

Several considerations may account for the differences in SASP factors identified through our approach compared to previous work. First, our approach did not rely on *in vitro*-based signatures to identify SnCs, which are known to poorly reflect senescence observed *in vivo*. In particular, *in vitro* assays induce senescence via artificial stressors that are not physiologically relevant to an *in vivo* environment. These stressors induce irreversible damage to the cell that contributes to the development of a pro-inflammatory SASP profile^69^, therefore skewing signatures derived from *in vitro* toward inflammatory cell types. Second, our study focused on cells expressing p16 and p16-derived signatures and did not focus on p21+p16-SnCs, which have been shown to correlate more strongly to inflammation *in vivo*^72–74^. Third, the SnCs observed in fibrosis may differ from those observed with inflammaging and inflammatory conditions.

Our study is the first to connect senotypes to distinct spatial niches in a fibrotic environment. Some previous studies have revealed that certain fibroblast SnCs *in vivo* are inflammatory^75,76^, while others may produce ECM and suppress inflammation^70,77^. Using integrated scRNA-seq and VisiumHD analysis, our approach accounted for those differences by mapping SnCs to different spatial microenvironments, offering insights into each senotype that could not be obtained through either modality alone. One distinct cellular niche (the “SnC Fibrotic Signaling” niche) contained myofibroblast SnCs alongside *Spp1*^hi^ macrophages, neutrophils, and blood vessels. Unexpectedly, this niche had a high cell density and regionally low ECM, suggesting that although these myofibroblast SnCs express fibrosis-associated ECM, they may instead stimulate fibrosis production in neighboring cells and participate in immunosuppression. Another significant niche contained immune signaling fibroblast SnCs alongside lymphocytes and antigen-presenting myeloid cells among loosely-organized ECM (the “SnC Immune Signaling & Tissue Remodeling” niche). The colocalization of these cells suggests that Immune Signaling fibroblast SnCs communicate with innate and adaptive immune cells through their SASP factors such as *Cxcl12* and *Saa3* to regulate immune cell recruitment, inflammation, and tissue remodeling. Additionally, the SnC Immune Signaling & Tissue Remodeling niche is often spatially adjacent to the SnC Fibrotic Signaling niche, suggesting potential cross-niche interactions that may orchestrate distinct but complementary processes in fibrosis. Finally, an observed niche containing Cartilage Developmment SnCs was embedded in highly-aligned mature ECM fibrils that were not associated with immune cells (the “SnC Cartilage Development” niche). The Cartilage Development niche occurred adjacent to each of the previous niches, suggesting that it may be a terminal ECM-rich niche that develops from fibrotic and immune signaling cues in the surrounding microenvironment. These observations are consistent with end-stage fibrotic matrix tissue deposition that has been observed in other fibrotic conditions^15^.

The fibrotic and immune signaling senotype niches resembled others that have recently been discovered in other fibrotic conditions. Studies have shown that distinct fibrotic and lymphoid-rich niches are present in idiopathic pulmonary fibrosis and interstitial lung disease, but not in healthy lung tissue^1–3,15^. The fibrotic niches were found adjacent to airway niches containing recruited/inflammatory macrophages and *Spp1*+ macrophages^1–3,15^, similar to the subpopulations observed at the implant interface in our model. The lymphoid-rich niches were adjacent but not overlapping with the fibrotic niches and included high expression of inflammatory SASP gene *Cxcl12*^1,2^. Similar fibroblast-containing niches have been observed in various cancers. One niche associated with myCAF exhibits increased ECM production and immunosuppression, contributing to decreased lymphocyte infiltration and poor prognosis^8,70,78^. On the other hand, a niche associated with iCAF exhibits enriched lymphocytes such as activated T cells, contributing to improved immunotherapy response and good prognosis^78,79^. The similarity of these lung- and cancer-related niches to those observed in our study suggests that similar senotypes may be participating in their divergent responses. However, future work is needed to understand the precise function of senotypes in such niches.

Our data also unveiled a timepoint-dependent emergence of fibroblast senotypes following muscle and lung injuries. Myofibroblast SnCs, including those high in *Spp1* expression, emerged during the acute healing stages and only persisted in injury environments with chronic inflammation and fibrosis. By contrast, Cartilage Development SnCs only emerged at later timepoints, during periods of active fibrosis. Recent studies have found that fibroblasts follow a differentiation trajectory after an inflammatory or stress-related stimulus, where they develop a fibrotic phenotype via a transient inflammatory state^51,53^. In the public bleomycin-induced lung injury dataset^51^, it was shown that Cthrc1+ fibrotic fibroblasts—including *Spp1*-high and *Fmod*-high subpopulations—originate from alveolar fibroblasts and differentiate along this trajectory. However, we found that these distinct *Spp1*-high and *Fmod*-high fibrotic populations remained present at the later stages of lung and muscle injuries, and appeared in different spatial niches in our muscle fibrosis model. Therefore, it remains unclear whether *Spp1*-high SnCs ultimately differentiate into a cartilage-like phenotype, or if they die off as inflammatory and fibrotic cues are dampened. Future studies using lineage tracing of *Spp1*+ fibroblasts could better elucidate the fate of such cells following fibrosis.

Furthermore, our observations provide insight into the development of primary and secondary SnCs *in vivo*. As described initially from *in vitro* studies, primary SnCs consist of cells that become senescent as a result of a stress stimulus, such as oncogene- or chemotherapy-induced senescence^80,81^. These SnCs express SASP and juxtacrine factors—such as those in the TGFβ, VEGF, and Notch signaling pathways—that can induce senescence in neighboring cells^80,82,83^. By contrast, secondary SnCs consist of cells that become senescent following exposure to SASP produced by a primary SnCs. While secondary SnCs undergo cell cycle arrest and upregulate SASP, such as fibrillar collagens, they may not induce senescence as effectively in other cells compared to primary SnCs^80,84^.

Although our data did not directly differentiate primary and secondary senescence, we observed spatial, temporal, and transcriptomic patterns consistent with these classifications. Myofibroblast SnCs emerged during the acute inflammatory stages in our model and were associated with “hotspots” of senescence at the later-stage fibrotic timepoint—suggesting that they have the potential to spread senescence to neighboring cells over time. In addition, myofibroblast SnCs express factors known to be linked to the induction of senescence in surrounding cells, including *Tgfb1*, *Vegfc*, *Notch1*, and *Jag1*^80,82,83^. However, myofibroblast SnCs also expressed fibrillar collagens—a finding consistent with the secondary senescence phenotype^84^. This suggests that myofibroblast SnCs could serve either as primary or secondary SnCs *in vivo*. Comparatively, Cartilage Development fibroblast SnCs emerged at later timepoints and in ECM-dense regions adjacent to the myofibroblasts or immune signaling senotype niches. These cells solely expressed SASP factors associated with ECM production and remodeling. They also expressed higher levels of *Tgfbr1* and *Tgfbr2*—receptors known to contribute to secondary senescence induction^80^. This suggests that Cartilage Development fibroblast SnCs serve as secondary SnCs.

We also observed timepoint-dependent development of the ECM Degradation fibroblast senotype—a subset of the Immune Signaling fibroblast senotype—in muscle injuries treated with a pro-fibrotic (PCL) or pro-regenerative material (dECM). These materials induce different healing outcomes: PCL promotes chronic inflammation and fibrosis, while dECM supports tissue repair. Despite these divergent processes, the ECM Degradation fibroblast senotype was observed in both conditions at the later timepoint, when both fibrosis and tissue remodeling occur. These data suggest that this senotype may be involved in processes beyond fibrosis, some of which could be beneficial to tissue healing. This concept aligns with published literature showing that fibroblasts expressing ECM degradation proteins, including MMPs and cathepsins, may serve either anti- or pro-fibrotic functions depending on context^85,86^. Future studies are needed to determine the function of this senotype and its utility for therapeutic targeting in fibrosis.

Our study is also the first to profile the perivascular SnC subtype in fibrotic conditions, highlighting its potential importance in regulating neovascularization and fibrosis in response to inflammatory cues. While the involvement of pericytes in fibrosis is well documented^87^, studies are conflicted on whether pericytes contribute to bulk fibrotic ECM production^5,59,60,88^, and the role of senescence in pericyte function during fibrosis remains unknown. By combining transcriptomics, multiplex IF, and pericyte-lineage SnC deletion models, we were able to elucidate the phenotype of perivascular SnCs and their functional role in a fibrotic context. Perivascular SnCs were located within and between the fibrotic and immune signaling senotype niches, and they upregulated pathways related to inflammatory responses, cell migration, and ECM remodeling. These observations are consistent with an activated phenotype that may influence vascular development and fibrosis^5,17,59,60^. By combining the pericyte lineage Ng2-CreER™ and p16-LOX-ATTAC models, we discovered that pericyte-lineage SnCs suppress the formation of mature vasculature and may modulate fibrosis production. These findings align with recent reports showing that some subsets of pericytes regulate microvascular remodeling and attenuate fibrosis in cardiac and lung injury models^18,59,60^. While the extent to which senescent pericytes are involved in such conditions remains unclear, our findings demonstrate the potential benefits of perivascular SnCs. Our findings also highlight the need to further study perivascular SnCs’ roles in fibrosis and other diseases to inform senolytic development.

Our study has several limitations. First, senotype-specific SASP profiles were determined through transcriptomics, which may not correlate with protein expression in each senotype. Single cell-level protein detection of SASP factors *in vivo* remains challenging due to the lack of working IHC antibodies for many SASP proteins and the inaccessibility of single cell proteomics techniques. While we confirmed the protein expression of many SASP factors in the bulk tissue, we cannot definitively link these proteins to specific cell types. Second, we lack Cre models that can target the specific fibroblast senotypes discovered in our study. The development of such models will be important to determine the functional relevance of fibroblast senotypes in fibrosis and other age-associated diseases. Third, our results were focused on studying senotypes with p16 expression and a p16-associated senescence signature (SenSig), so they may not have captured p21-associated senotypes, such as the p21+p16-myeloid cells observed in our model. Further studies are needed to determine the phenotypes and functions of such SnCs.

## Materials and Methods

### Experimental animals

All animal procedures performed in this study were approved by the Johns Hopkins Animal Care and Use Committee under protocols MO21M80 and MO24M66. Mice were housed in the Johns Hopkins Research Animal Resources central animal facilities. We obtained the following strains for this study: C57BL/6 (Jackson Laboratories, Strain No. 000664), C57BL/6 from NIA rodent colony, and Ng2-CreER^TM^ (Jackson Laboratories, Strain No. 008538^89^). The p16-LOX-ATTAC strain^61^ was developed and donated by Dr. Sundeep Khosla from the Mayo Clinic. Heterozygous/homozygous Ng2-CreER^TM^ and homozygous p16-LOX-ATTAC were crossed to generate a hybrid Ng2-CreER^TM^;p16-LOX-ATTAC strain to enable selective pericyte-lineage senescent cell elimination. Ear snips <3 mm in diameter were collected and submitted to Transnetyx for genotyping. We used 6-8-week-old female mice for all experiments involving the C57BL/6 strain. For pericyte-lineage elimination studies, we used 11-week-old male mice heterozygous for the Ng2-CreER^TM^ and p16-LOX-ATTAC transgenes.

### Clinical samples

Human keloid tissue samples were obtained from deidentified surgical discards of patients undergoing keloid removal surgery at the Johns Hopkins Hospital. Samples were obtained under a Johns Hopkins approved IRB exemption IRB00088842. Tissues were fixed in 10% formalin for at least 96 hours prior to processing. Tissue embedding and sectioning was performed by the Johns Hopkins University Oncology Tissue and Imaging Service SKCCC core. Tissues were sectioned at a 4 μm thickness using a microtome. Tissue blocks and sections were stored at room temperature.

### VML injury and biomaterial implantation procedure

Bilaterial VML procedures were performed as previously described^25^. All surgical tools were autoclaved prior to surgery. PCL particles (<600 μm diameter, MW: 50000 kDa, Polysciences 25090 or Ingevity CAPA® 6506) were sterilized under an ultraviolet lamp within a biosafety cabinet for 2-4 hours and were stored in sterile tubes for <1 week prior to surgical procedures. Mice were anesthetized using 1.5-3% isoflurane and administered 1 mg/kg Buprenorphine SR (Wedgewood Pharmacy) by intraperitoneal injection to provide preoperative analgesia. Prior to incision, hair was removed from the site of incision using clippers, and the underlying skin was sterilized using 70% ethanol. To begin the surgical procedure, a ∼1 cm longitudinal skin incision was made over the quadriceps muscle, and the underlying fascia was gently cut away from the muscle. A ∼3×3×3 mm^3^ defect was made in the quadriceps muscle using sterile microdissection scissors. Special care was given to avoid tearing the quadriceps tendon during the defect creation. PCL particles were implanted into the defect space using a sterile surgical scoop (Moria Surgical 1121b, 1 scoop per defect, scoop dimensions: 8 mm diameter, 1 mm depth). The skin incision was closed using sterile 5-0 nylon sutures. The same procedure was repeated on the other quadriceps muscle. Following the procedure, mice were kept on a heat pad until they were fully awake and ambulatory. Mice were routinely monitored starting 24 hours after surgery until they completely recovered.

### Ng2+ cell lineage SnC elimination

Heterozygous 11-week-old male Ng2-CreER^TM^;p16-LOX-ATTAC mice were used to study the effects of eliminating pericyte (Ng2+ cell) lineage SnC deletion on vascularization and fibrosis in the murine VML-PCL injury model. To enable selective targeting of pericyte-lineage SnCs, Cre recombination was induced in Ng2+ cells using tamoxifen administration. Tamoxifen was administered by intraperitoneal (IP) injection (70 mg/kg) daily for five consecutive days. To prepare tamoxifen solution, tamoxifen powder (MilliporeSigma, Cat. No. T5648) in corn oil (MilliporeSigma, Cat. No. C8267) at a concentration of 14 mg/mL overnight at 37°C on an orbital shaker. The solution was stored at 4°C and protected from light. Aliquots of the solution were pre-heated for at least 10 minutes at 37°C prior to the injections. The tamoxifen solution was used within seven days following preparation. At Day 14 following the first injection of tamoxifen, bilateral VML injury and PCL implantation surgeries were performed as described earlier. To eliminate pericyte-lineage SnCs, AP20187 (Takara Bio, Cat. No. 632622) was administered twice weekly by IP injection (10 mg/kg) starting at Day 4 following surgery. Fresh AP20187 (2 mg/mL AP20187, 4% v/v Ethanol, 10% v/v PEG-400, 2% Tween-20 in distilled water) and vehicle solutions (4% v/v Ethanol, 10% v/v PEG-400, 2% Tween-20 in distilled water) were prepared immediately prior to injections using the manufacturer’s protocol for IP injections in mice. Mice were euthanized after 6 weeks following the VML injury and biomaterial implant procedure, and quadriceps tissues were harvested as described earlier.

### Murine tissue processing and sectioning

Mice were euthanizing using an overdose of isoflurane following by cervical dislocation. Following euthanasia, quadriceps muscles containing the PCL implants were carefully removed using dissecting scissors and fixed in 10% formalin for 48 hours with gentle agitation at room temperature. Tissues were rinsed twice with PBS for at least 5 min each and stored in 70% ethanol at room temperature for less than 4 weeks. To prepare tissues for paraffin embedding, tissues were dehydrated using a graded series of ethanol (1 hour each in 80%, 95%, 100%, 100%) and cleared using xylenes (2 x 45 min). Tissues were incubated in paraffin at 60°C for 12-24 hours. Paraffin was exchanged 4-6 times to remove residual xylenes from the tissues. Immediately before embedding, tissues were placed on a cooling block and bisected along the cross section of the mid-point of the defect. Tissues were embedded in paraffin molds with each cross section facing downward and were chilled on a cooling block for at least 15 min. Tissue blocks were stored at room temperature prior to sectioning.

Tissue sectioning for histology, multiplex immunofluorescence staining, and VisiumHD was performed by the Johns Hopkins University Oncology Tissue and Imaging Service SKCCC core. Tissues were sectioned at a 7 μm thickness using a microtome. Sections for VisiumHD were processed according to the manufacturer’s protocol under RNAse-free conditions. Sections were stored either at room temperature (histology, multiplex immunofluorescence) or -20°C (VisiumHD).

### Histological staining and brightfield imaging

H&E and Masson’s Trichrome staining was performed by the Johns Hopkins University Oncology Tissue and Imaging Service SKCCC core using standard protocols. All sections within the same experiment were stained at the same time to ensure consistency across samples. Brightfield imaging was performed using the Hamamatsu NanoZoomer (Model C12000-02). Slides were scanned at 40X magnification. Images were exported as .JPG files using NDPView 2. No linear or non-linear adjustments were made to the images displayed in the figures.

### Multiplex immunofluorescence staining and imaging

Multiplex immunofluorescence staining was performed using the Opal system from Quanterix (previously Akoya Biosciences), as previously described^25^. Steps were performed at room temperature unless indicated otherwise. The list of primary antibodies and dilution factors used in this study is provided in **Supplemental Table 1**. Slides were baked in an oven at 60°C to promote tissue adherence and remove excess paraffin. Slides were then deparaffinized and rehydrated using the following incubations: 3 x 3 min in xylenes, 1 min each in graded ethanol series (100% x 2, 95%, 80%, 70% ethanol), and 3 x 3 min in MilliQ water. Antigen retrieval was performed in pre-heated 1X AR6 buffer (Quanterix, Cat. No. AR6001KT) for 15 min in a vegetable steamer. Slides were cooled for 15 min in a room temperature water bath, rinsed once in MilliQ water, and incubated in 3% hydrogen peroxide for 15 min on a shaker to block endogenous peroxidases. Blocking was performed for 30 min by incubating the tissues in a solution of 10% bovine serum albumin (BSA) and 0.05% Tween-20 in 1X DPBS. Immediately following blocking, tissues were stained with the first primary antibody for 30 min. All primary antibodies were diluted in blocking buffer prior to staining. Following incubation in primary antibody, tissues were washed for 3 x 3 min in 1X TBST (Cell Signaling, Cat. No. 9997s) and stained in secondary antibodies raised against the host species of the primary antibody (**Supplemental Table 1**). Then, tissues were washed for 3 x 3 min in 1X TBST, incubated in Opal dye diluted in 1X Manual Amplification Diluent for 10 min, and washed again for 3 x 3 min in 1X TBST. Opal dyes and dilutions used in this study are listed in **Supplemental Table 1**. To start the next round of staining, tissues were incubated in pre-heated 1X AR6 buffer for 15 minutes in a vegetable steamer, cooled in a room temperature water bath for 15 minutes, and rinsed in MilliQ water. Steps for blocking, primary antibody, secondary antibody, and Opal incubations were repeated for each round of staining (one primary antibody per round, 2-3 rounds total). The 1X AR6 incubation and cooling steps were performed after each round of staining to strip the antibodies from the tissue. For experiments requiring three rounds of staining, tissues were stored in 1X TBST at 4°C overnight between staining rounds. After the final round of Opal dye incubation, tissues were washed in TBST for 3 x 3 min, rinsed in MilliQ water, and incubated in 1X Spectral DAPI (Quanterix, Cat. No. FP1490) for 5 min. Tissues were then rinsed twice in MilliQ water and mounted using DAKO Fluorescent Mounting Medium (Agilent, Cat. No. S302380-2) and 1.5 thickness coverslips. Slides were dried for >12 hours at 4°C before imaging.

**Supplemental Table 1.** List of antibodies and Opal dyes used in this study.

| Antibody or Opal Dye | Source | Dilution & Incubation Time |
| --- | --- | --- |
| Primary Antibodies |  |  |
| p16 (mouse tissues) | Abcam, Cat. No. ab211542 | 1:1000, 30 min |
| p21 | Abcam, Cat. No. ab188224 | 1:1000, 30 min |
| Ki67 (D3B5) | Cell Signaling Technology, Cat. No. 12202 | 1:200, 30 min |
| CD31 | Abcam, Cat. No. ab182981 | 1:2000, 30 min |
| CD11b | Abcam, Cat. No. ab133357 | 1:1000, 30 min |
| MMP3 | Abcam, Cat. No. ab52915 | 1:50, 30 min |
| SPP1/Osteopontin | R&D systems, Cat. No. MAB808-SP | 1:200, 30 min |
| PDGFRβ | Cell Signaling, Cat. No. 3169 | 1:100, 30 min |
| CD146 | Abcam, Cat. No. ab75769 | 1:500, 30 min |
| p16 (human tissues) | CINTEC-Roche, Cat. No. 705-4793 | 1:5, 15 min |
| CD31 (human tissues) | Abcam, Cat. No. ab76533 | 1:100, 30 min |
| Secondary Antibodies |  |  |
| Rabbit-on-rodent HRP-Polymer (mouse tissues) | Biocare Medical, Cat. No. RMR622H | 30 min |
| MACH3 Mouse HRP-Polymer Detection (human tissues, mouse antibody detection) | Biocare Medical, Cat. No. M3M530 | 10 min in Probe<br>Wash 3x3 min in 1X TBST<br>10 min in HRP Polymer |
| MACH4 Universal HRP-Polymer (human tissues, rabbit antibody detection) | Biocare Medical, Cat. No. M4U534 | 10 min in Probe<br>Wash 3x3 min in 1X TBST<br>10 min in HRP Polymer |
| Opal Dyes |  |  |
| Opal 520 | Quanterix, Cat. No. FP1487001KT | 1:100, 10 min |
| Opal 570 | Quanterix, Cat. No. FP1488001KT | 1:150, 10 min |
| Opal 650 | Quanterix, Cat. No. FP1496001KT | 1:150, 10 min |

Tissues were imaged using the Zeiss Axio Observer.Z1/7. Images were acquired as a z-stack (3 μm interval) using the 10X objective and with the apotome engaged. Tile scans were performed using a 10% overlap between tiles. For each experiment, exposure times for each dye were set based on a ∼30% shift in the histogram, and exposure times were held constant between samples. Images were stitched and flattened to a maximum intensity projection using ZenBlue software (Carl Zeiss). Linear adjustments to the images included in the figures were performed using ZenBlue. No non-linear adjustments were made to the images unless specified in the figure captions.

### Image analysis

Cell abundance and marker colocalization were quantified using HALO v3.6 with the HighPlex FL v3.2 module (Indica Labs). The annotation tool was used to draw regions of interest around the PCL implant for measurements. Skeletal muscle and adipose tissue adjacent to the implant were excluded from analysis. Regions of tissue folding within the PCL implant were also excluded. Cell nuclei were segmented based on DAPI nuclear staining. The cytoplasm of each cell was defined as the area within a 1 μm radius from the edge of the cell nucleus. Manually defined thresholds were set for each nuclear and cytoplasmic stain to delineate cells positive and negative for each marker. Cell abundance and colocalization measurements were exported from the software and analyzed using Microsoft Excel and GraphPad Prism 11.

Vessel area analysis was performed using QuPath-0.6.0. Regions of interest were drawn around the PCL implant, as described above. A custom script employing the OpenCV Pixel Classifier plugin was used to detect blood vessels. Vessels were detected using a manually defined threshold and a gaussian filter (sigma = 0.3) of the CD31 stain. The threshold was held constant across all images within an experiment. CD31+ fragments <20 μm^2^ or >900,000,000 μm^2^ were excluded from the analysis. The resulting vessel annotation layer was split into one annotation per vessel fragment to enable individual vessel area measurements. Measurements were exported and analyzed in Microsoft Excel and GraphPad Prism 11.

### Tissue proteomics

Proteomics data were derived from a published dataset of the muscle fibrotic tissue implants (MassIVE ID Number: MSV000098194). Detailed methods used to collect and process tissues, conduct liquid chromatography-tandem mass spectrometry (HPLC-MS/MS), and process the data are described in a previous publication^20^. Briefly, PCL implants from VML-injured quadriceps muscle of C57/BL6 female mice were harvested at Week 6 following implantation, cut away from the surrounding muscle tissue, flash frozen on dry ice, and frozen at -80 °C until further processing. Tissues were thawed, homogenized, digested, and desalted to isolate peptides for HPLC-MS/MS analysis. HPLC-MS/MS was performed using an Evosep One liquid chromatography system (Evosep) coupled to a timsTOF HT mass spectrometer (Bruker). The resulting data were processed in Spectronaut (v19.7.250203.62635) using the directDIA workflow. Peptides were matched to proteins using the *Mus musculus* reference proteome (UniProtKB, reviewed entries only), accessed on 3/27/2025. Proteins with ≥2 unique peptides were included in the analysis. Protein abundance values were based on the peak areas of extracted ion chromatograms of 3 – 6 MS2 fragment ions with local normalization and q-value sparse data filtering applied and iRT profiling selected. From these data, SASP proteins were identified, as described in Ext. Data 3, and plotted on a heatmap using -log(10) of the raw protein abundance value.

### Spatial transcriptomics (VisiumHD)

Spatial transcriptomics was performed using the VisiumHD Mouse Transcriptome 6.5 mm kit with the v2.0 probe set (10x Genomics, Cat. No. 1000676). Tissue sections were prepared as described earlier and stored at -20°C until processing. Prior to VisiumHD, each tissue block was assessed for QC using four 7 μm-thick tissue curls per block. RNA extraction and QC measurements were performed by Psomagen (Rockville, MD, USA). Both tissue blocks used for VisiumHD analysis passed the manufacturer’s QC recommendation (DV200>30%). H&E staining, brightfield imaging, CytAssist probe capture and probe extension, library construction, and sequencing were performed by Psomagen according to the manufacturer’s protocols.

### scRNA-seq analysis of CD45-enriched dataset

#### Myeloid cell subclustering

The scRNA-seq data was preprocessed, clustered and annotated as described previously^20^. The dataset was subset to the clusters containing myeloid cells (neutrophils, monocytes, macrophages, dendritic cells) and reprocessed using Seurat v4.4.0 according to previously described methods^20^. To avoid potential artifacts associated with tissue digestion, cell stress-related genes previously linked to tissue digestion were omitted during the re-clustering analysis^90^. The filtered counts were log normalized and scaled using the ScaleData function in Seurat. Principle component analysis (PCA) was performed on the top 2000 variable features using the scaled data, and an elbow plot was generated to determine the optimal number of principle components (10 PCs). A shared nearest neighbor graph was generated using the first 10 PCs via the FindNeighbors function. Clustering was then performed using the Louvain algorithm via the FindNeighbors function. Cluster marker genes were identified using the FindAllMarkers function with default parameters (Wilcoxon Rank Sum test) on the non-filtered RNA counts. Cluster resolution (0.325) was selected based on the stability of the clusters, as visualized using Clustree v0.5.1, and the presence of distinct markers that defined each cluster. Clusters were annotated using their top differentially expressed genes.

#### SenSig-projectR transfer learning

Transfer learning with SenSig^25^ was conducted using the projectR^23,24^ v1.22.0 package, as previously published^23^. First, the scRNA-seq object and SenSig were preprocessed prior to input into projectR. In the scRNA-seq object, genes were filtered to include only those present in SenSig (FDR<0.05). Normalized expression values of those genes were then scaled using z-scoring for input into projectR. SenSig was re-processed to enable projectR transfer learning (GSE199864)^25^. The source bulk RNA-seq data (GSE199864)^25^ were processed for differential expression analysis using the standard pipeline in edgeR v4.6.3^91^. Differentially expressed genes were filtered to only include those with FDR<0.05. The F-statistic for each of those genes was converted to a t-statistic using the following formula: sign(logFC) × 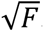. The t-statistic was then converted to a standardized z-statistic using the zscoreT function in edgeR. projectR transfer learning was performed by projecting the scRNA-seq dataset into SenSig using the scRNA-seq z-score scaled gene expression and SenSig z-statistic values as inputs. The outputs of projectR included a projection weight and p-value for each cell. Cells with a positive projection weight and p-value <0.01 were considered senescent.

#### Differential gene expression and gene set enrichment analysis (GSEA)

Differential gene expression analysis between cells classified as SnC and not SnC in designated clusters was performed using a Wilcoxon Rank Sum test via the Seurat v5.4.0 FindMarkers function using default parameters. Secreted ligands, matrisome, and cell surfaceome genes were annotated using the CellPhoneDB v4.1.0^29^, SEPDB^30^, MatrisomeDB^31^, and SURFY^32^ databases. Genes with an adjusted p-value <0.05 were considered differentially expressed.

GSEA between SnCs and not SnCs in designated clusters was performed using fgsea v1.32.4 package. Ranked gene lists were obtained by re-running the FindMarkers function in Seurat, as described above. The following parameters were set in FindMarkers to obtain differential expression results for all genes without filtering: logfc.threshold = 0, min.pct = 0, min.cells.group = 1, min.cells.feature = 1. Genes were ranked by the product of their log_2_FC and -log_10_(p-value) values. Gene sets from the Hallmark^92^, Reactome^93^, and GOBP^94,95^ databases were obtained from MSigDB^96^ (msigdbr v10.0.2) and combined for the analysis. GSEA was conducted via the fsgea function using the ranked gene list and list of combined pathways as inputs. Pathways with an adjusted p-value <0.05 were considered significant.

#### TF activity inference and differential TF activation analysis

TF activity was predicted using the pySCENIC v0.12.1 package^46^ based on methods and code previously published^20^. Differential TF activation between SnC and not SnC in designated clusters was performed using a Wilcoxon Rank Sum test on the median area under the curve (AUC) scores of each TF. A Benjamini-Hochberg correction was applied to account for multiple comparisons. TFs with an adjusted p-value <0.05 were considered significantly activated in SnCs versus not SnCs.

#### Coordinated Gene Activity across Pattern Subsets (CoGAPS) analysis

CoGAPS, a non-negative matrix factorization algorithm, was conducted on the fibroblast clusters, as previously described^42^, to identify biological patterns associated with fibroblast senotypes. Fibroblast clusters—including the two progenitor fibroblasts, activated fibroblasts, *Lrrc15+* myofibroblasts, cycling fibroblasts, and cartilage-like fibroblasts—were computationally isolated and input into CoGAPS for analysis. Low variance genes (counts with standard deviation ≤0.1) were removed from the dataset. A log transformation was applied to the counts of the remaining genes.

CoGAPS v3.20.0 was run on the log transformed data using the following parameters: 50000 iterations, sparse optimization enabled, and genome-wide parallelization enabled. A total of 10 CoGAPS runs were made using a different pattern number (*k*) as input. The optimal CoGAPS dimensions were selected by evaluating the drop in mean Chi-squared value across the different dimensions, as described in the Main text. Pattern marker genes were determined using the patternMarkers function in CoGAPS using the “cut” threshold parameter. This approach selects marker genes for each pattern based on whether each gene is less significant to at least one other pattern. Patterns were manually annotated based on the association of their marker genes with known biological processes and/or fibroblast cell subpopulations.

To determine the correlation between CoGAPS patterns and TF activity, a Spearman correlation between the CoGAPS pattern (sampleFactors) matrix and pySCENIC TF AUC matrix was computed using the Hmisc v5.2.5 package. Correlations with a p-value <0.05 were considered significant.

To determine the specificity of CoGAPS patterns to SnCs, the values of the pattern (sampleFactors) matrix were compared between SnC and not SnC within each cluster. Each row (pattern) in the sampleFactors matrix was compared between SnC and not SnC using a t-test with Benjamini-Hochberg correction.

#### Comparison of senescence signatures

Multiple senescence signatures were evaluated to determine their similarity to the senotypes discovered from SenSig-projectR and CoGAPS analyses. GSEA was conducted as described earlier using the senescence signatures as input into fgsea. Senescence gene sets with an adjusted p-value <0.1 were considered significant. The scores of select senescence signatures were computed on the entire scRNA-seq dataset using the Seurat v5.4.0 AddModuleScore function. To evaluate the relationship between SenMayo and CoGAPS pattern weights in fibroblasts, the fibroblasts clusters were computationally isolated, and the SenMayo module score was computed using the AddModuleScore function. The Spearman correlation calculation was performed between the SenMayo module score and CoGAPS pattern weights (extracted from sampleFactors matrix) for each CoGAPS pattern at the *k* = 12 and *k =* 18 dimensions.

### VisiumHD analysis

#### Data preprocessing

Raw data were processed by Psomagen (Rockville, MD, USA) using the manufacturer’s protocol (10x Genomics). Space Ranger v3.1.0 was used to align the raw FASTQ files to the mouse mm10-2020-A (GENCODE vM23) reference genome, align the fiducials, and count UMIs in each bin.

#### Data processing and analysis using Seurat

Processed data were loaded into Seurat v5.4.0^27^ using the 8 μm bin output files from Cell Ranger and normalized for downstream analyses. The data were subsetted to 50000 bins using the SketchData function in Seurat. The “leverage score” method was used to enable capture of rare subpopulations. Clustering was performed on the sketched data according to the standard Seurat pipeline, as described earlier in the scRNA-seq methods, using 50 PCs and a cluster resolution of 0.5. The clustering results were projected into the full dataset using the default parameters in the ProjectData function. Cluster marker genes were identified using the default parameters of the FindAllMarkers function. Clusters were annotated by comparing their marker genes to those of the CD45-enriched scRNA-seq clusters and to established canonical markers for different cell types. Clusters that spatially overlapped with blank regions (e.g. regions with no tissue) were annotated as “junk” and omitted from further analysis.

#### projectR transfer learning with SenSig and CoGAPS patterns

Transfer learning via projectR v1.22.0^23^ was conducted to project the VisiumHD data into SenSig and the CoGAPS patterns, as described earlier. Clusters containing skeletal muscle cells—which were minimally present in the fibrotic implant tissue—were omitted from these analyses to enhance computational efficiency. For transfer learning with SenSig, normalized data were filtered to include genes only present in SenSig. Additional filtering was then performed to exclude low variance genes (standard deviation ≤ 0.5). Transfer learning via projectR was performed using the filtered genes and SenSig z-statistic as the input. Z-score scaling was performed on the filtered genes prior to input into projectR. The z-scored dataset set was split into smaller chunks before running projectR for better memory efficiency. Bins with a positive projection weight and p-value less than 0.01 were considered senescent.

Transfer learning with CoGAPS was conducted using a similar chunking process as with SenSig. The amplitude (featureLoadings) matrix from the *k* = 12 matrix was filtered to exclude the two patterns annotated as “noise.” Genes were filtered to include only those present in the featureLoadings matrix. Normalized expression of the filtered genes and the filtered featureLoadings matrix were used as input into projectR. The projection results were used for data visualization and analysis.

#### Data processing and analysis using STAPLE

Processed data from Space Ranger were analyzed using the STAPLE v2.2.6-g0f8ee7a^97^ pipeline to perform spatial co-occurrence and neighborhood analysis. Prior to running STAPLE, cell type deconvolution was performed using RCTD as implemented in spacexr v2.2.1^57^. To enable cell type deconvolution, the CD45-enriched scRNA-seq dataset was reformatted to annotate VisiumHD bins based on cell class, including fibroblasts (all fibroblast clusters), vascular cells (ECs, perivascular cells), lymphatic ECs, T/NK cells, and myeloid cells (Neutrophils 1 & 2, Monocytes/Early Macrophages, *Spp1*^hi^ Macrophages 1, *Mrc1*+ Macrophages). Skeletal muscle cells, mast cells, satellite cells, and *Spp1*^hi^ macrophages 2 were omitted due to their low cell numbers and/or lower quality relative to the other clusters. This reprocessed scRNA-seq object was input to RCTD as the reference along with the VisiumHD data to annotate the 8 μm bins. Bins with labels assigned as “Unknown” were omitted from the analysis. To obtain more information about cell subtypes, the bins annotated as myeloid cells and fibroblasts were computationally isolated and subclustered using a similar procedure as the described in the Seurat analysis. Clusters were annotated using a similar process as described in the Seurat analysis. The RCTD and the Space Ranger output files were used as the input for STAPLE.

The data were ingested into the STAPLE pipeline as AnnData v0.12.6^98^ objects, and the Squidpy v1.6.6.dev24+g32789aefd^58^ module was used to analyze pairwise spatial co-occurrence and cell type neighborhoods. This analysis was performed both for the lower resolution (without myeloid cell and fibroblast subclusters) and the higher resolution cell type labels.

### SENgEAR analysis of public scRNA-seq datasets

SENgEAR was used to query the senotypes discovered from our fibrosis model in other injury and fibrotic conditions. We obtained public datasets for muscle injury with different biomaterial treatments^25^ (GSE175890), bleomycin-induced lung fibrosis^51^ (GSE210341), and human keloid^63^ (GSE163973). Datasets were uploaded to SENgEAR using the standard gEAR workflow^19^. The SenSig (z-statistic for genes with FDR<0.05) and CoGAPS featureLoading matrices were uploaded to gEAR to enable transfer learning. Datasets were projected into SenSig and CoGAPS patterns using the least squares optimization algorithm in the SENgEAR “Projection Tool.” For the projections involving SenSig, genes were scaled using the “Z-score normalize gene expression” option. Negative coefficient weights were set to zero for visualization. Violin and UMAP visualizations for each dataset were designed using the “Single-gene Displays” workflow.

### Data visualization

scRNA-seq data were visualized using the Seurat (v5.0.0-v5.4.0)^27^, ggplot2 (v3.5.0-v4.0.2)^99^, Tidyverse (v2.0.0)^100^, and ComplexHeatmap (v2.20.0)^101,102^ packages in R. Select bar plots and heatmaps were generated using GraphPad Prism v10-v11. VisiumHD data were visualized using Seurat (v5.0.0-v5.4.0)^27^, ggplot2 (v4.0.2)^99^, Tidyverse (v2.0.0)^100^, and Squidpy (v1.6.6.dev24+g32789aefd)^58^. scRNA-seq data analyzed in SENgEAR were visualized as described earlier. Quantitative microscopy data were visualized using GraphPad Prism v10-v11.

### Statistics

Statistics for scRNA-seq and VisiumHD analyses are described in the Main text and in the earlier Methods sections. All other statistics were conducted using GraphPad Prism v10-v11. Statistical tests used in GraphPad Prism are described in the figure captions. Significance for all tests was defined as a p-value or adjusted p-value <0.05 unless indicated otherwis.

## Data Availability

Spatial transcriptomics data generated for this study will be made publicly available for download on GEO prior to publication. The CD45-enriched scRNA-seq dataset used in this paper was previously published^20^ and can be downloaded under GEO accession number GSE306253. The proteomics dataset was previously published^20^ and can be downloaded at the Mass Spectrometry Interactive Virtual Environment (MassIVE) repository using the following link: https://massive.ucsd.edu/ProteoSAFe/dataset.jsp?task=6f0be9baa689478b8989ab9ec098f20b (MassIVE ID Number: MSV000098194; ProteomeXchange ID: PXD065015). Other publicly available datasets used for this study were downloaded under GEO accession numbers GSE175890 (muscle VML with different timepoints and treatments^25^), GSE210341 (bleomycin-induced murine lung fibrosis^51^), and GSE163973 (human normal and keloid scars^63^). All scRNA-seq datasets and senescence signatures (SenSig, CoGAPS patterns) in this work will be made publicly available on the interactive SENgEAR web portal to enable custom visualizations and analyses. Raw microscopy data will be made publicly available on the Biostudies repository prior to publication. All other data are available in the manuscript or supplemental materials.

## Code Availability

Codes used for processing the CD45-enriched dataset, including clustering and SCENIC analysis, are available on the Elisseeff lab Github at the following link: https://github.com/Elisseeff-Lab/cd45enriched_fibrosis_scRNAseq. Code for all other analyses will be made publicly accessible prior to publication on the Elisseeff lab Github repository: https://github.com/Elisseeff-Lab. The SENgEAR web portal and associated datasets may be accessed at this link: https://senescence.umgear.org/

## Author Contributions

A.N.R., S.N., E.J.F., and J.H.E conceptualized and designed the studies.

A.N.R. and A.C. performed experiments and interpreted results. P.S.D., M.B., J.C.M., and A.R. assisted with experimental work.

S.N. and E.J.F. developed computational pipelines.

S.N. and A.N.R. and performed computational analyses and interpreted results.

R.S.A., D.L., J.O., S.N., A.A.M., O.R.W., and E.J.F. developed the SENgEAR web platform and uploaded datasets and senotype signatures to the platform.

K. K. and F.H.Y. assisted with scRNA-seq computational analyses.

M.B. and A.N.R. managed the transgenic murine colonies.

D.P.R., C.D.K., and B.S. designed, performed, and interpreted the tissue proteomics experiments.

K.O.B. provided deidentified clinical tissue specimens.

A.N.R. and J.H.E. wrote the manuscript.

J.H.E. and E.J.F. supervised the study.

## Declaration of Interests

J.H.E holds equity in Unity Biotechnology and Aegeria Soft Tissue and is a consultant for Tessara. E.J.F has been on the Scientific Advisory Board of Resistance Bio / Viosera Therapeutics and was a paid consultant to Mestag Therapeutics and Piper Sandler, she is on the scientific advisory board for the V Foundation and has a familial relationship to the founder of PushCART Therapeutics.

## Acknowledgements

We would like to thank Dr. Sundeep Khosla and Jennifer Rowsey for providing the p16-LOX-ATTAC transgenic mouse strain and advising on colony management and Michelle Giglio for supporting analysis on data uploading. We thank the Johns Hopkins University Oncology Tissue and Imaging Service SKCCC core for performing histological sectioning and staining, Johns Hopkins University Research Animal Resources for managing the animal facilities, Johns Hopkins University Tumor Microenvironment Core for providing imaging and image analysis equipment, C M Cherry consulting for performing the initial processing of the CD45-enriched scRNA-seq dataset (GSE306253), and Psomagen for performing VisiumHD. Select schematics were created using Biorender.

## Funding

This study was supported by NIH grants 1U54AG079779 (J.H.E., E.J.F.), R01HD111243 (J.H.E.), R01AG082965 (J.H.E., E.J.F.), R01EB028796 (J.H.E.), DP1AR076959 (J.H.E.), K99AG081564 (J.C.M.), DGE1746891 (AR), and U54 AG075932 (B.S.). This study was also supported by the Bloomberg∼Kimmel Institute (J.H.E.) and the University of Maryland, Baltimore through a grant from the Pivot Strategic Investment Initiative (E.J.F.). The content is solely the responsibility of the authors and does not necessarily represent the official views of the University of Maryland, Baltimore.

---

**Extended Data 1.**
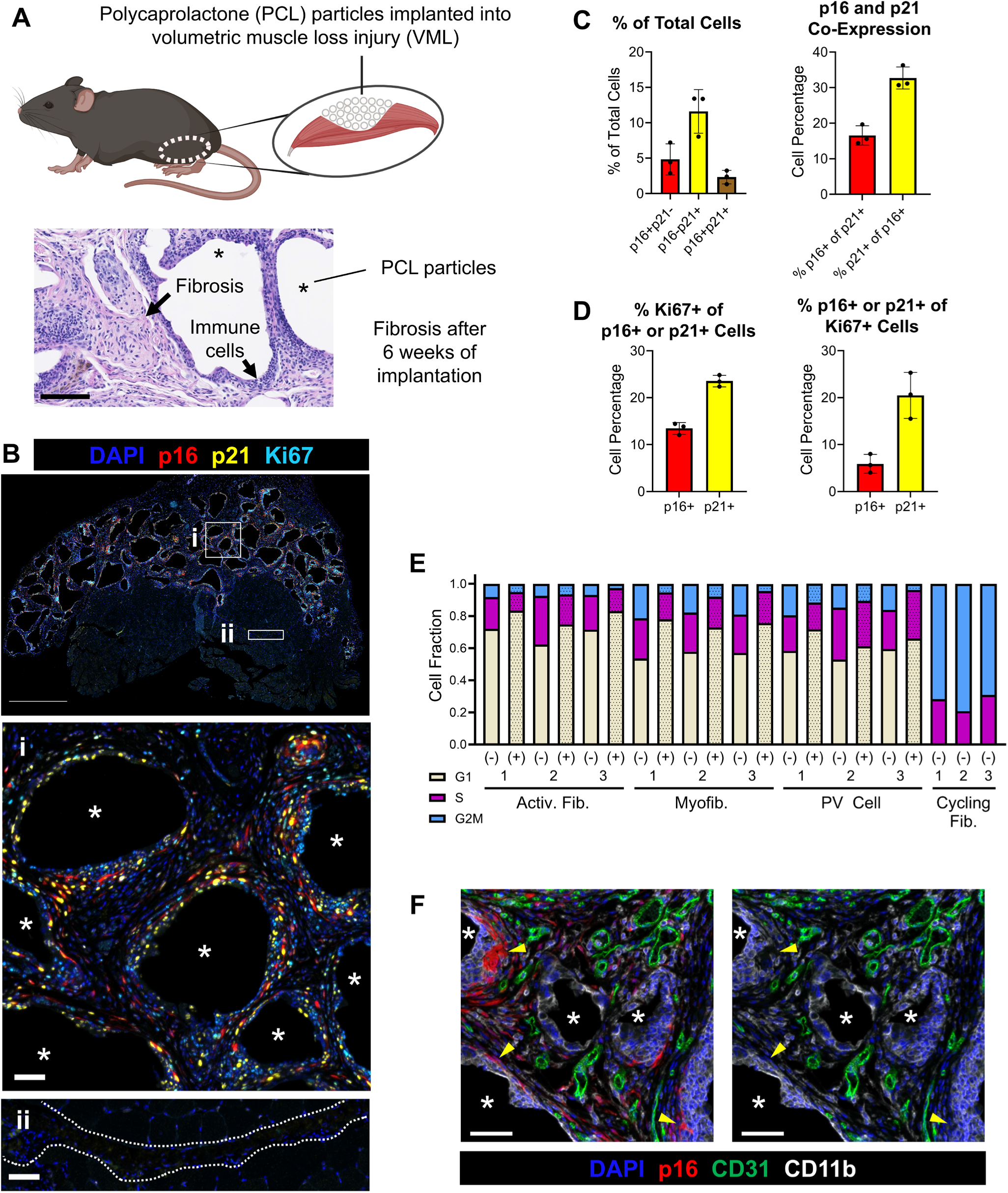
Biomaterial-induced fibrosis is enriched in cells positive for senescence markers. **A)** Schematic and H&E image depicting biomaterial-mediated fibrosis model. The model induces chronic inflammation and fibrosis around the implant particles by 6 weeks following injury and biomaterial implantation. Scale bar: 100 µm. *PCL (implant) particles. **B)** Immunofluorescence image of p16, p21, and Ki67 in the fibrotic implants. The bottom two images show the fibrotic implant region (i, *PCL particles) and the muscle fascia (ii, dotted outline). The brightness of the top image was increased by 40% using Microsoft Powerpoint to enable visualization of the full tissue. Scale bar: 1000 µm (top), 50 µm (bottom). **C)** Quantification of cells positive for p16 and/or p21. n=3 biological replicates **D)** Quantification of co-expression between Ki67 and p16 or p21. Data are mean ± standard deviation. n=3 biological replicates. **E)** Extension of plot shown in Figure 1D to show data from each biological replicate (labeled 1, 2, 3). **F)** Immunofluorescence images showing lack of expression of p16 in CD11b+ myeloid cells (yellow arrowheads). Scale bar: 50 µm. *PCL particles

**Extended Data 2.**
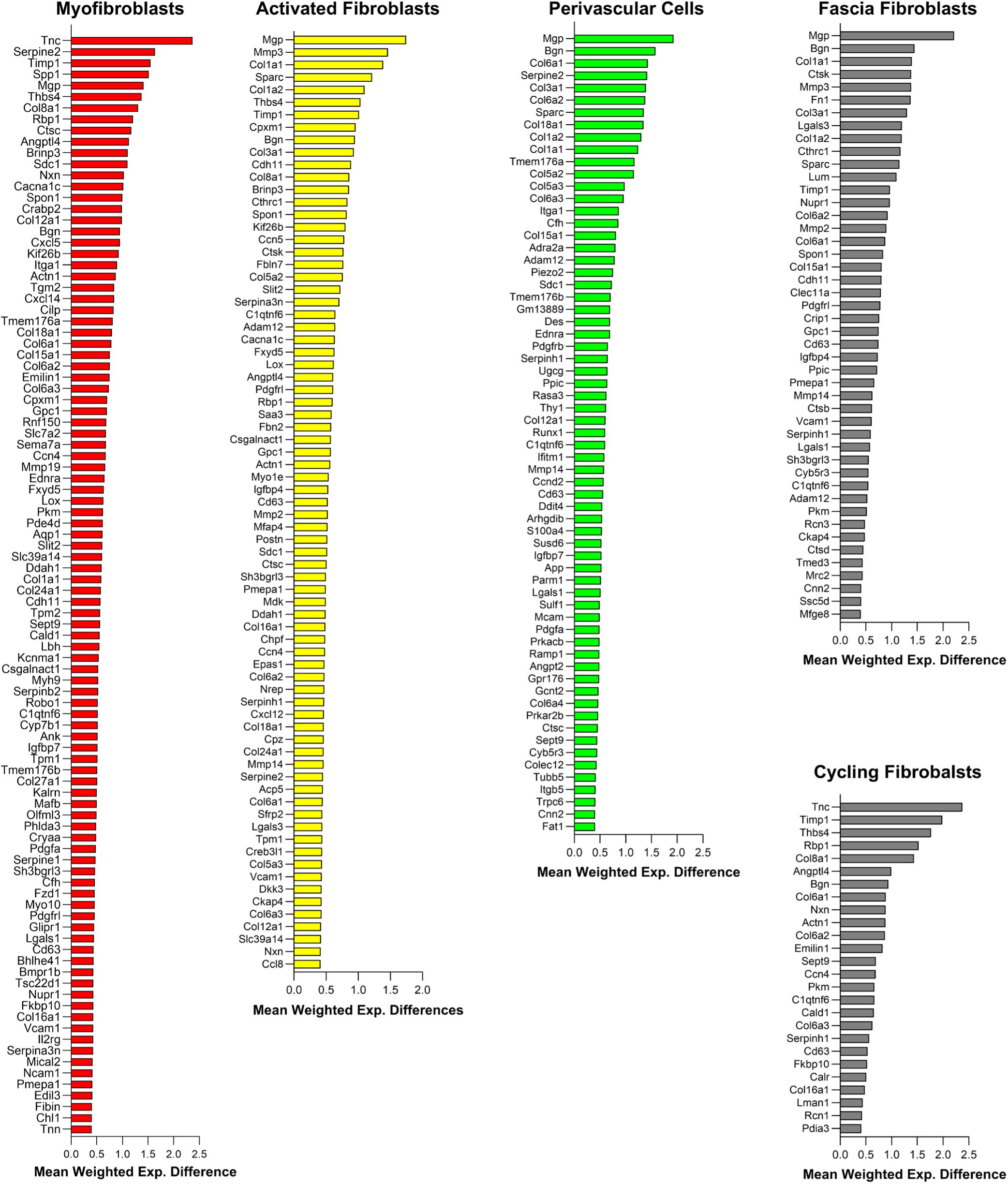
Projection driver genes in SenSig-high clusters. Driver genes were computed by comparing the weighted gene expression of SenSig genes classified as SnC and not SnC within each cluster. Genes with a weighted mean expression above 0.4 are shown in the plots.

**Extended Data 3.**
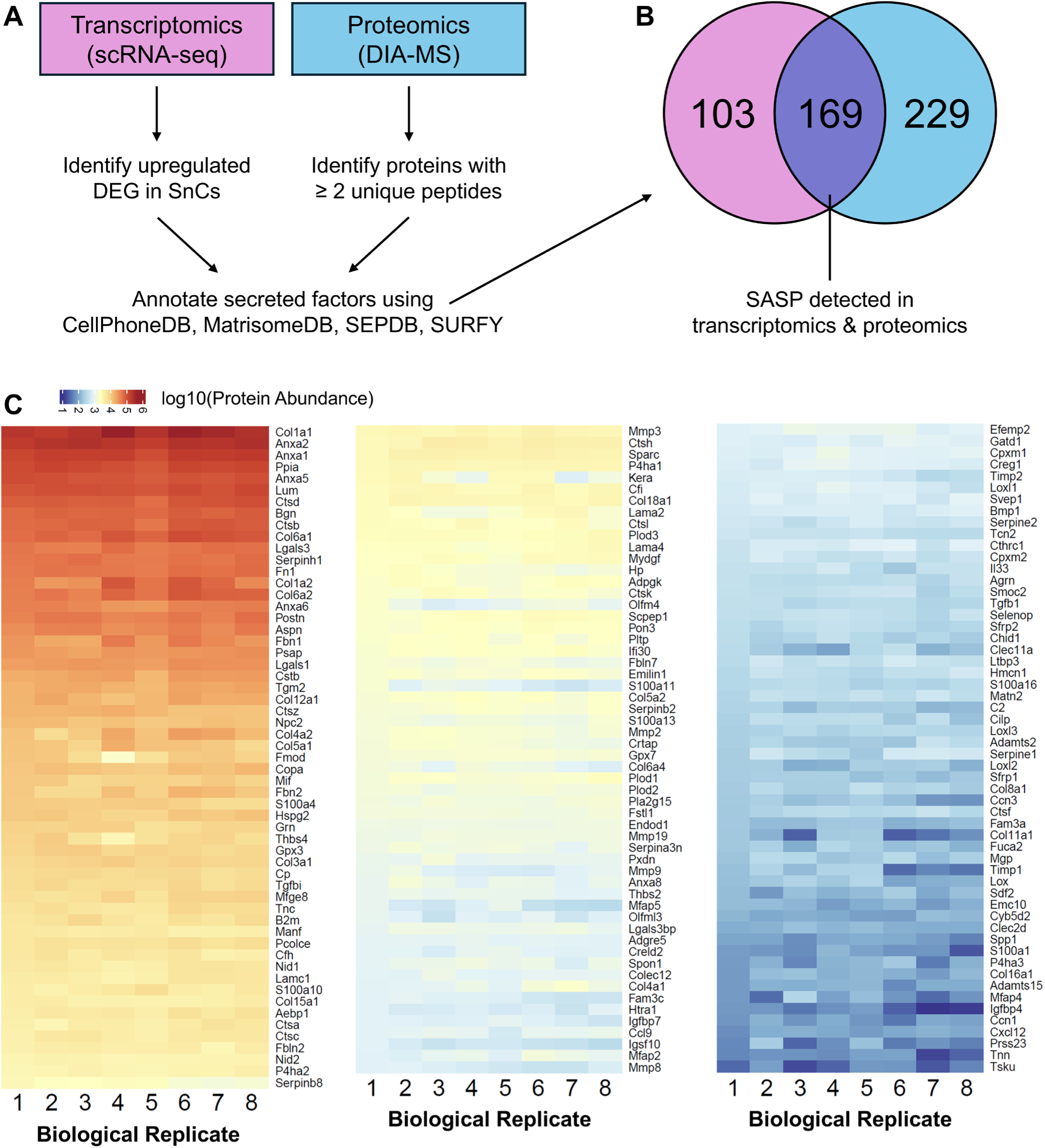
Proteomics confirms expression of predicted SASP factors in fibrotic tissue. **A)** Pipeline to identify secreted factors in scRNA-seq and proteomics data. Proteomics data were obtained using data-independent acquisition mass spectrometry (DIA-MS). Proteins with >=2 peptides detected in all samples were included in the analysis. Secreted factors were identified using the CellPhoneDB, MatrisomeDB, and SEPDB databases. Transmembrane proteins were omitted using the SURFY (cell surfaceome) database. **B)** Venn diagram displaying number of proteins detected in both the transcriptomics and proteomics analysis. **C)** Heatmap displaying log10 of protein abundance of SASP factors detected in the bulk fibrotic tissue. Each column represents a biological replicate (n=8).

**Extended Data 4.**
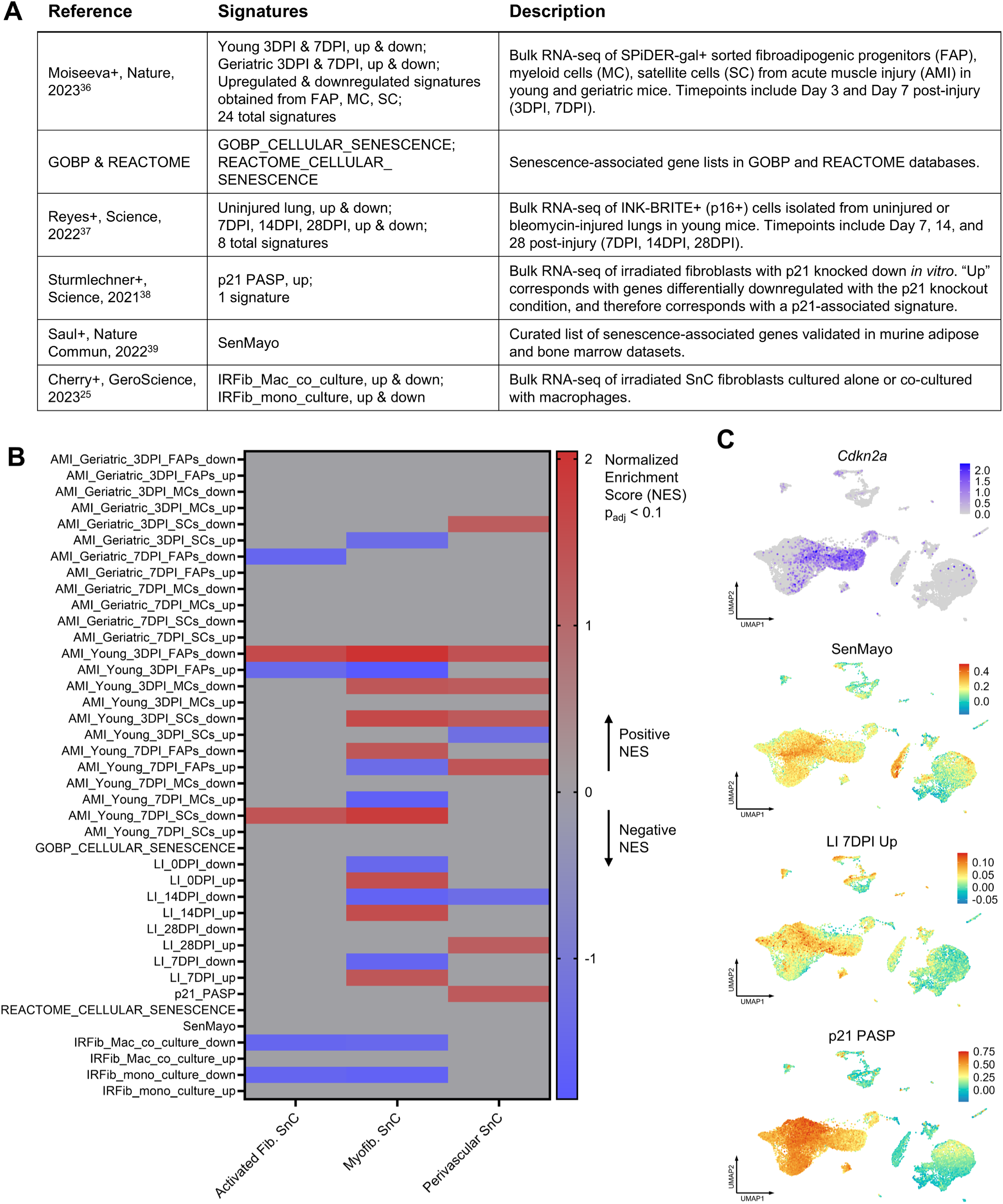
Comparison of fibroblast and perivascular senotypes with other published senescence signatures. **A)** Table describing murine senescence signatures used for the analysis. **B)** Gene set enrichment analysis using published senescence signatures. The normalized enrichment score is shown for results with an adjusted p-value (p_adj_) less than 0.1. Results with p_adj_ > 0.1 are displayed in gray. **C)** UMAP projection showing normalized expression of *Cdkn2a* and Seurat module score for select senescence signatures.

**Extended Data 5.**
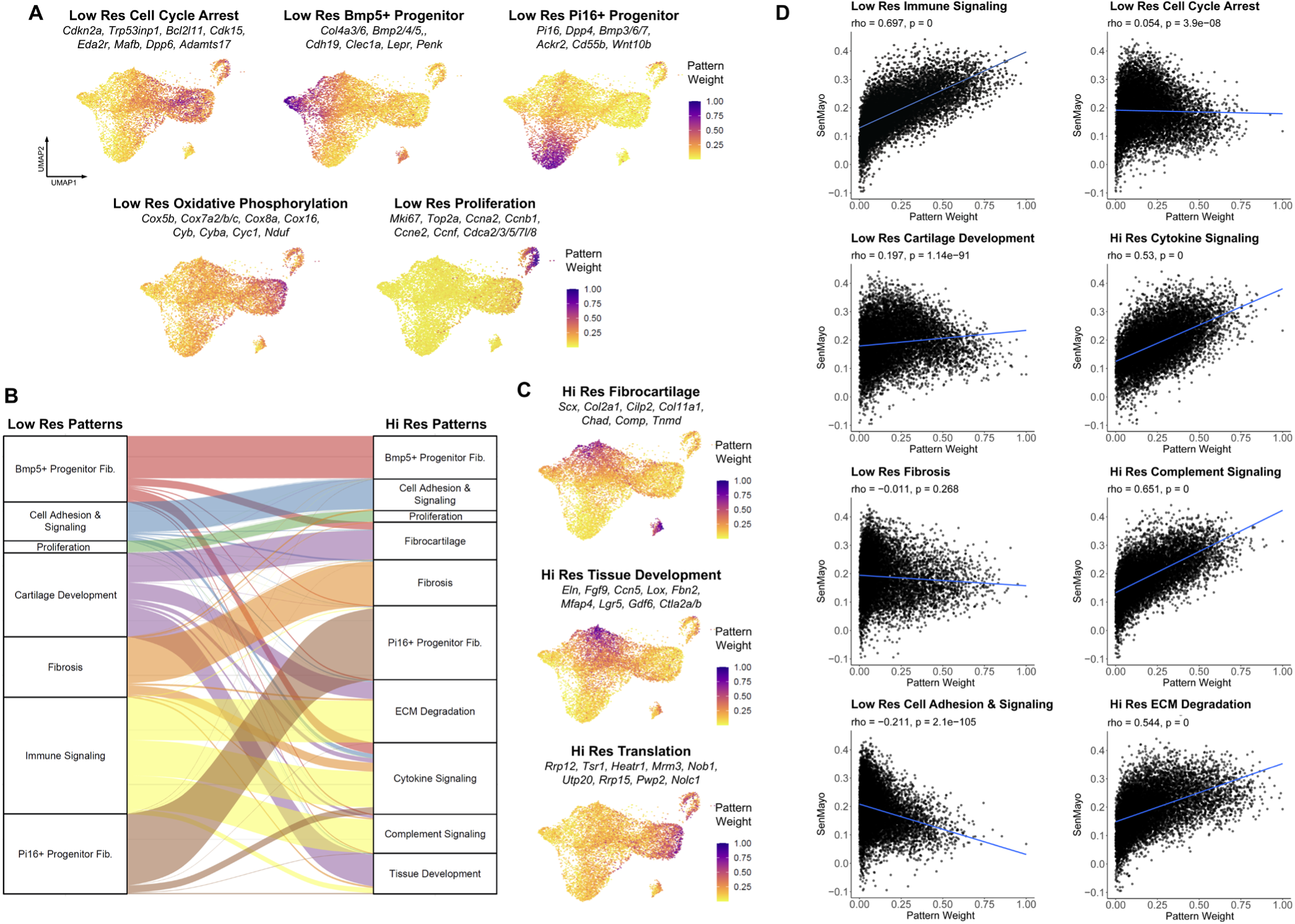
Additional CoGAPS patterns detected in fibroblasts. **A)** UMAP and marker genes of patterns detected at Low Res CoGAPS. **B)** Relationship between cell type-specific Low Res and Hi Res CoGAPS patterns representing distinct fibroblast subpopulations. Cells were assigned their most dominant pattern at the Low Res and Hi Res for the analysis. The number of cells per pattern is represented in the y-axis of the plot. **C)** UMAP and marker genes of additional patterns detected using Hi Res CoGAPS. Hi Res Fibrocartilage and Hi Res Tissue Development represent sub-patterns of Low Res Cartilage Development (**B**). **D)** Spearman correlation between senescence-associated Low Res and Hi Res CoGAPS pattern weights and SenMayo Seurat module score. The correlation coefficient and p-values are displayed above each plot.

**Extended Data 6.**
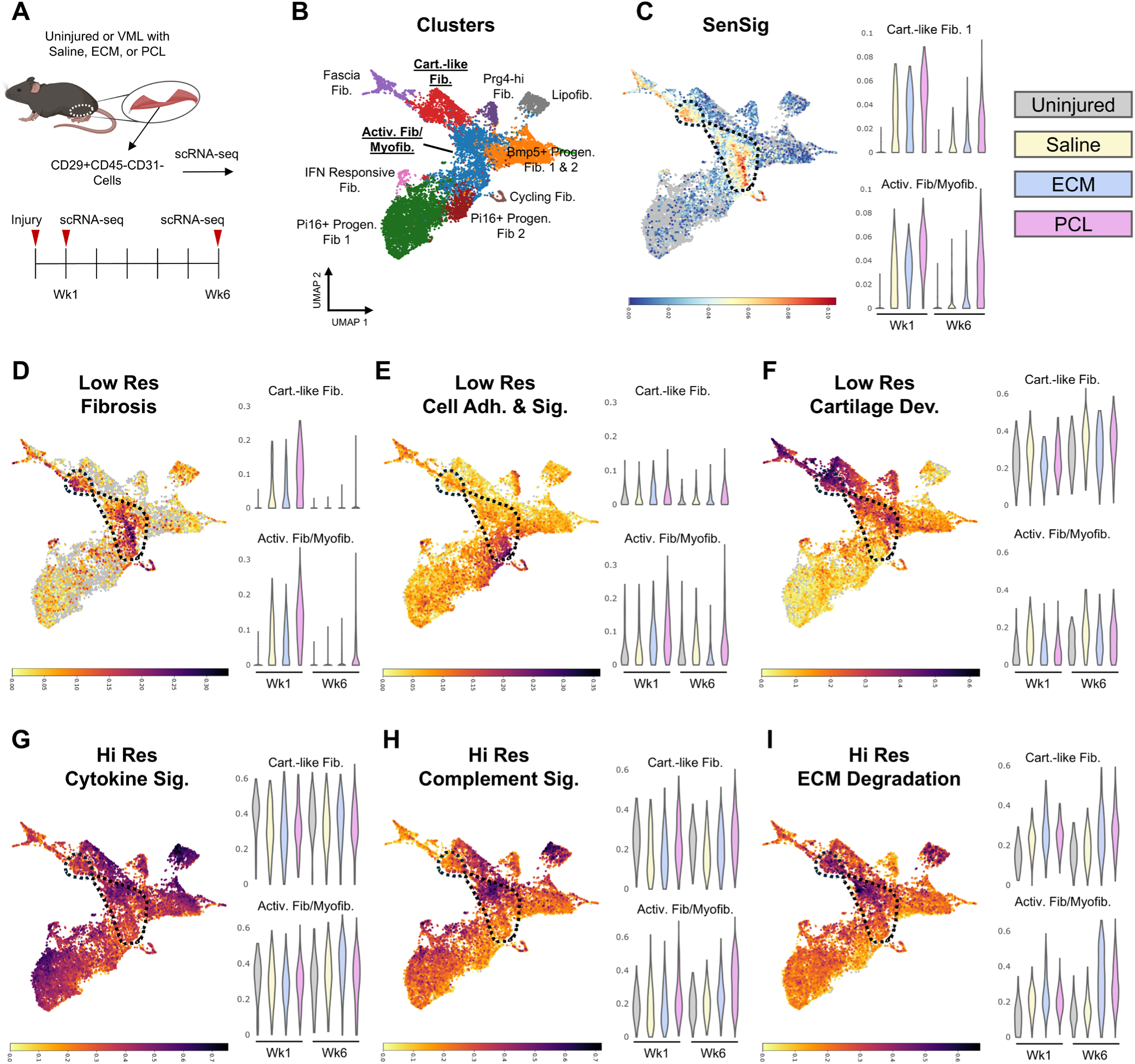
Fibroblast senotypes emerge across different stages of wound healing and fibrosis development. **A)** Experimental design used to obtain dataset in Cherry et al.^25^, GSE175890. CD29+CD45-CD31-cells were sorted from uninjured muscle or injured muscle treated with saline, decellularized ECM implant (ECM), and PCL and sequenced using Drop-Seq. Samples were collected at 1 and 6 weeks post-injury and treatment. Uninjured muscles were obtained from age-matched mice at each timepoint. **B)** UMAP of cell clusters. The clusters with the highest SenSig weights shown in **C** are underlined. **C-I)** Projection weight of SenSig (**C**) and CoGAPS patterns (**D-I**) obtained via transfer learning. The dotted outline shows cells with a high SenSig projection weight in the Activated Fibroblasts/Myofibroblasts and Cartilage-like Fibroblasts. Violin plots show projection weight in each experimental condition.

**Extended Data 7.**
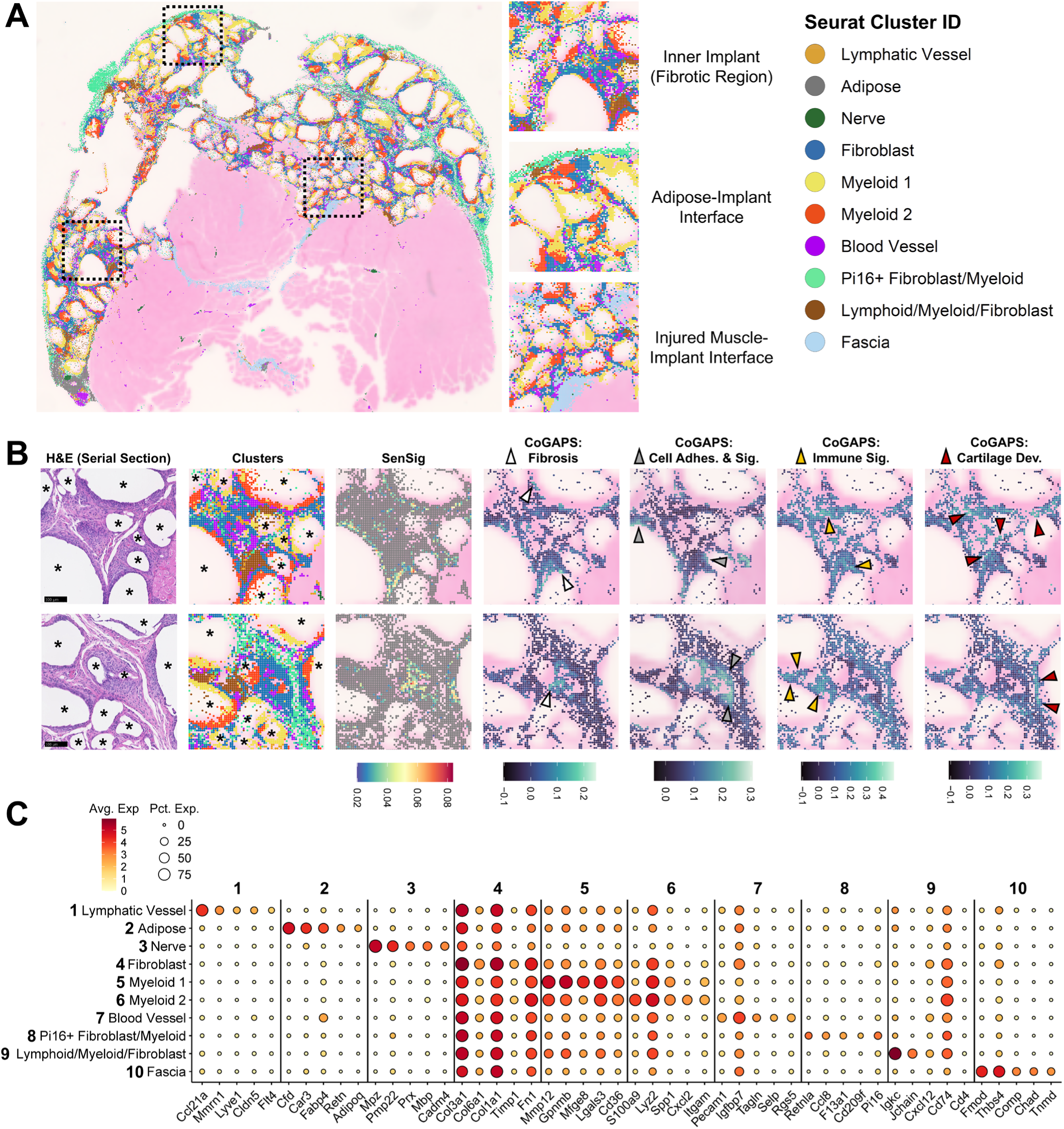
VisiumHD analysis of fibrotic muscle implant. **A)** Seurat-assigned clusters in fibrotic muscle implant. Insets show clusters present in the inner implant, adipose-implant interface, and muscle-implant interface. Clusters annotated as “Muscle,” “Red Blood Cell,” or “Junk” were omitted from the plot. **B)** Zoomed-in regions of the inner implant showing Seurat-assigned clusters, SenSig projection weight, and fibroblast CoGAPS pattern projection weight. CoGAPS patterns are only shown in bins assigned to fibroblast-containing clusters. Scale bar: 100 μm. **C)** Dot plot of select differentially expressed genes of each Seurat-assigned cluster. The color scale represents average scaled gene expression by cluster. The dot size corresponds to the percentage of bins within each cluster that express the gene.

**Extended Data 8.**
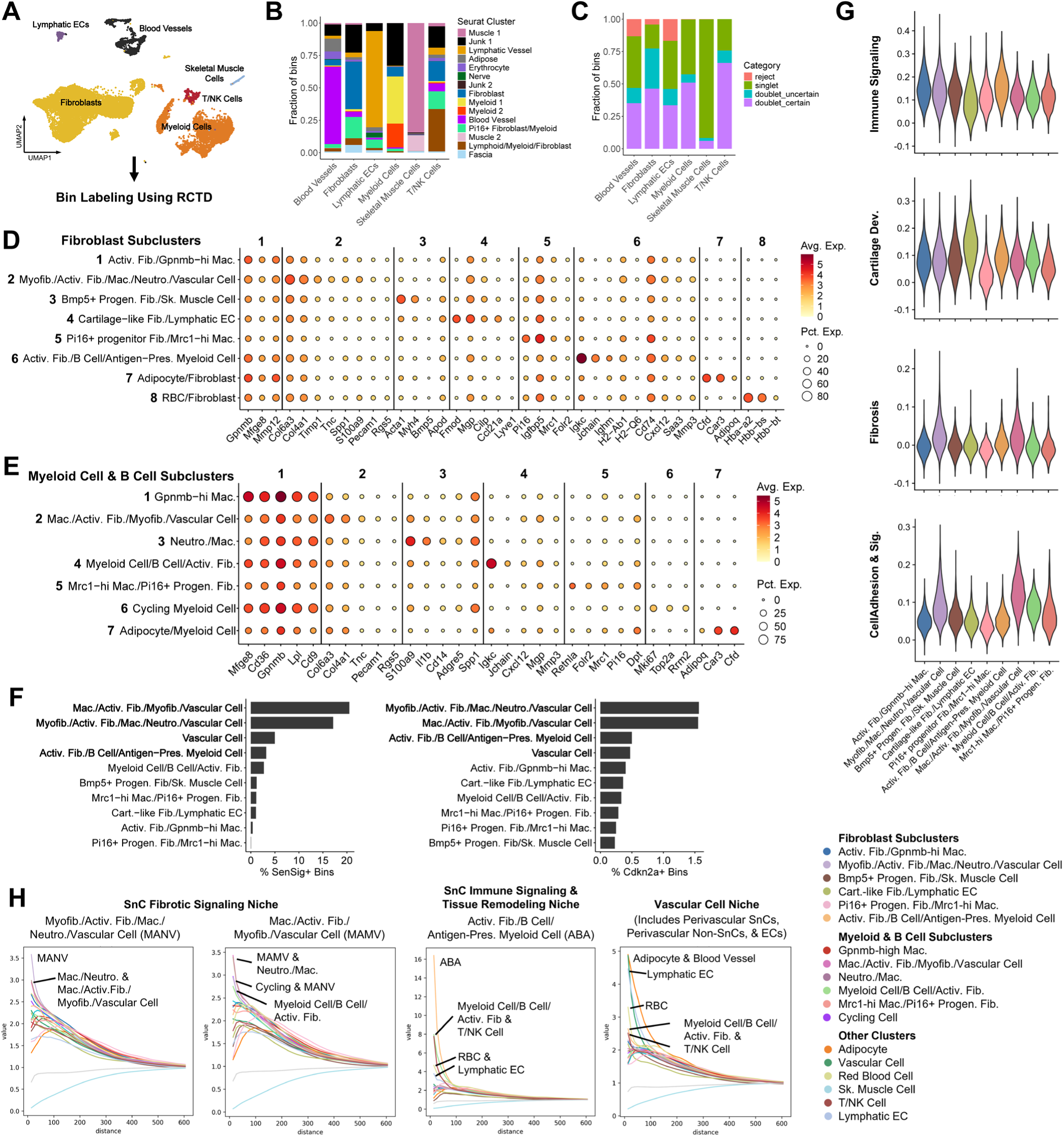
RCTD and Squidpy analysis reveals co-occurrence patterns for senotype niches. **A)** scRNA-seq labels used as input annotations for RCTD. **B)** Comparison of RCTD labels to Seurat clusters. **C)** Singlet and doublet rate in each RCTD-assigned bin label. **D-E)** Fibroblast (**D**) and Myeloid (**E**) subcluster markers. Plots show average normalized expression and percentage of cells expressing the marker for each subcluster. **F)** Bar plots showing percentage of SenSig+ and *Cdkn2a*+ bins in clusters containing fibroblasts or vascular cells. Clusters with the highest percentage of SenSig+ and *Cdkn2a*+ bins are shown in bold. **G)** Projection weight of fibroblast CoGAPS patterns in clusters containing fibroblasts. **H)** Squidpy co-occurrence analysis of clusters representing different SnC niches. The y-axis represents the co-occurrence probability value computed using the following equation: Probability(Cluster | Cluster of interest)/Probability(Cluster of Interest). The y-axis represents the co-occurrence probability within a given radius around a spot (x-axis). A higher probability value indicates a stronger chance of co-occurrence between two clusters at each radius.

**Extended Data 9.**
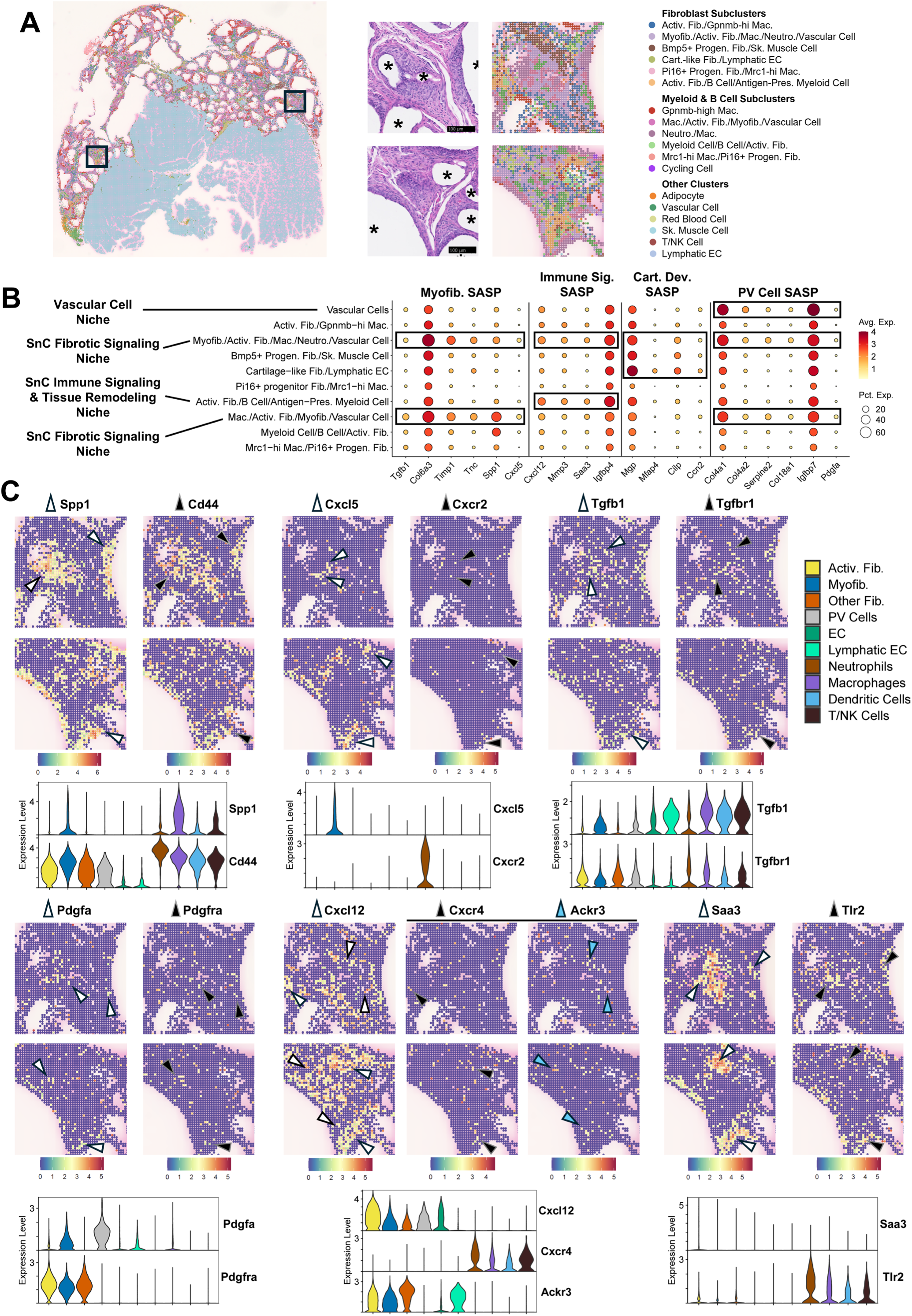
SASP ligand and receptor expression in senotype niches. **A)** Spatial plot of RCTD clusters and corresponding H&E images of select regions in the fibrotic implant. Asterisks (*) label regions of the PCL particle implant. Scale bar: 100 µm **B)** Senotype-specific SASP factor expression in RCTD clusters containing fibroblasts or vascular cells. Clusters containing high levels of SASP factor expression for each senotype are outlined in black. Clusters representing SnC niches are labeled in bold text on the left. **C)** Spatial plots showing expression of myofibroblast, immune signaling fibroblast, and perivascular SnC SASP ligands and their corresponding receptors in the regions shown in **A**. The color scale of each spatial plot represents normalized gene expression. Arrowheads point to regions of co-expression between the SASP ligands and receptors. Violin plots display normalized expression of SASP factors and receptors in the reference scRNA-seq object. The legend for each cell type included in the violin plots is shown on the right.

**Extended Data 10.**
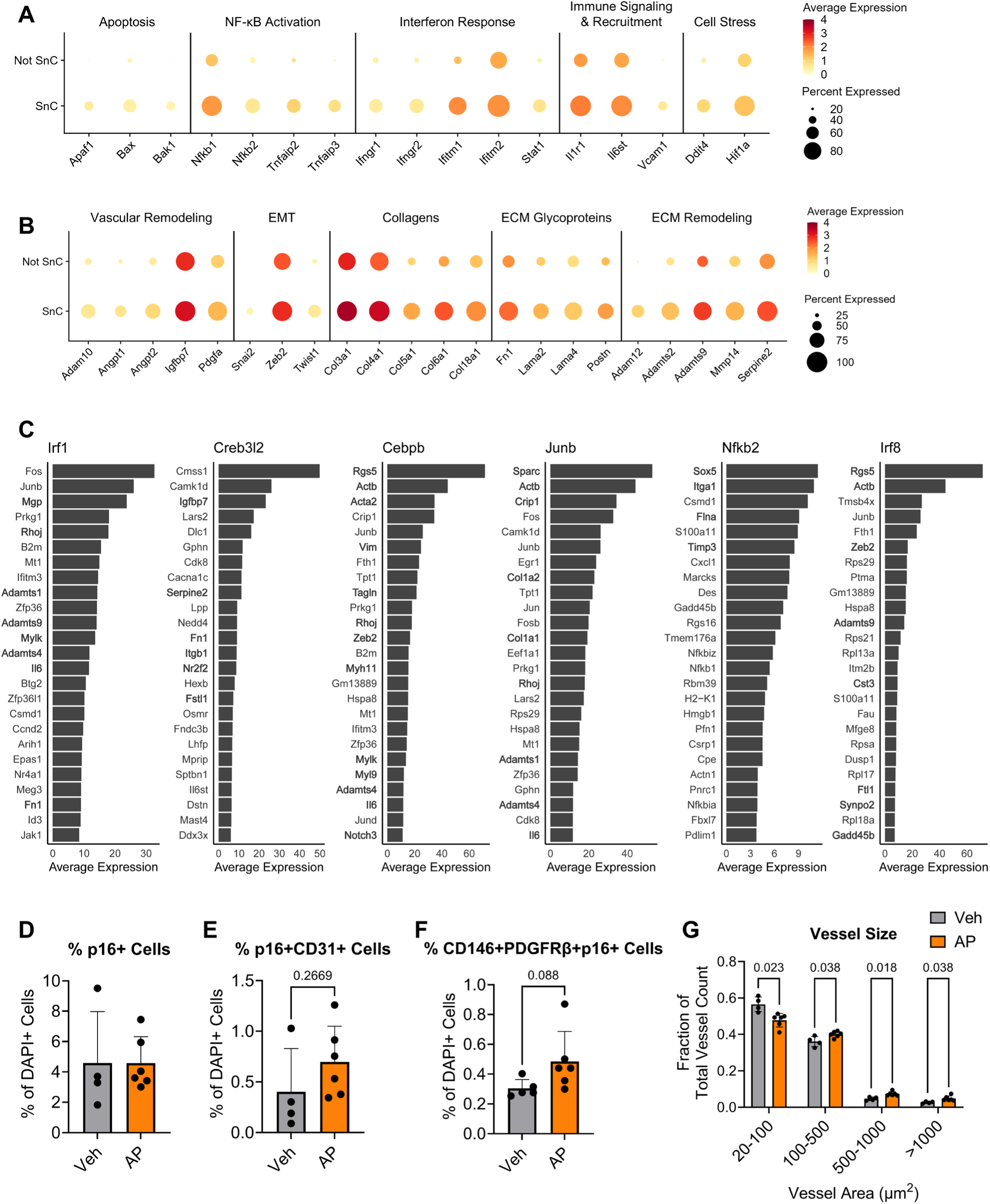
Perivascular SnCs upregulate genes associated with inflammatory responses and ECM remodeling. **A-B)** Average normalized expression of differentially upregulated genes related to apoptosis and cellular stress (**A**) and vascular and ECM remodeling (**B**) in perivascular SnCs. **C)** Top 25 genes in regulons related to cytokine responses and cellular stress. Downstream genes related to EMT, vascular remodeling, and ECM remodeling are shown in bold. Gene expression represents average expression within perivascular SnCs. **D-F)** Quantification of p16+, p16+CD31+, and p16+CD146+PDGFRβ+ cell percentage of total cells with (AP) and without (Veh) pericyte-lineage p16+ cell depletion. Two-tailed t-test. n=4-6 biological replicates. **G)** Fraction of vessels within each size bin (area). Two-tailed t-test with Holm-Šídák method for multiple comparisons. Data are mean ± standard deviation. n=4-6 biological replicates.

**Extended Data 11.**
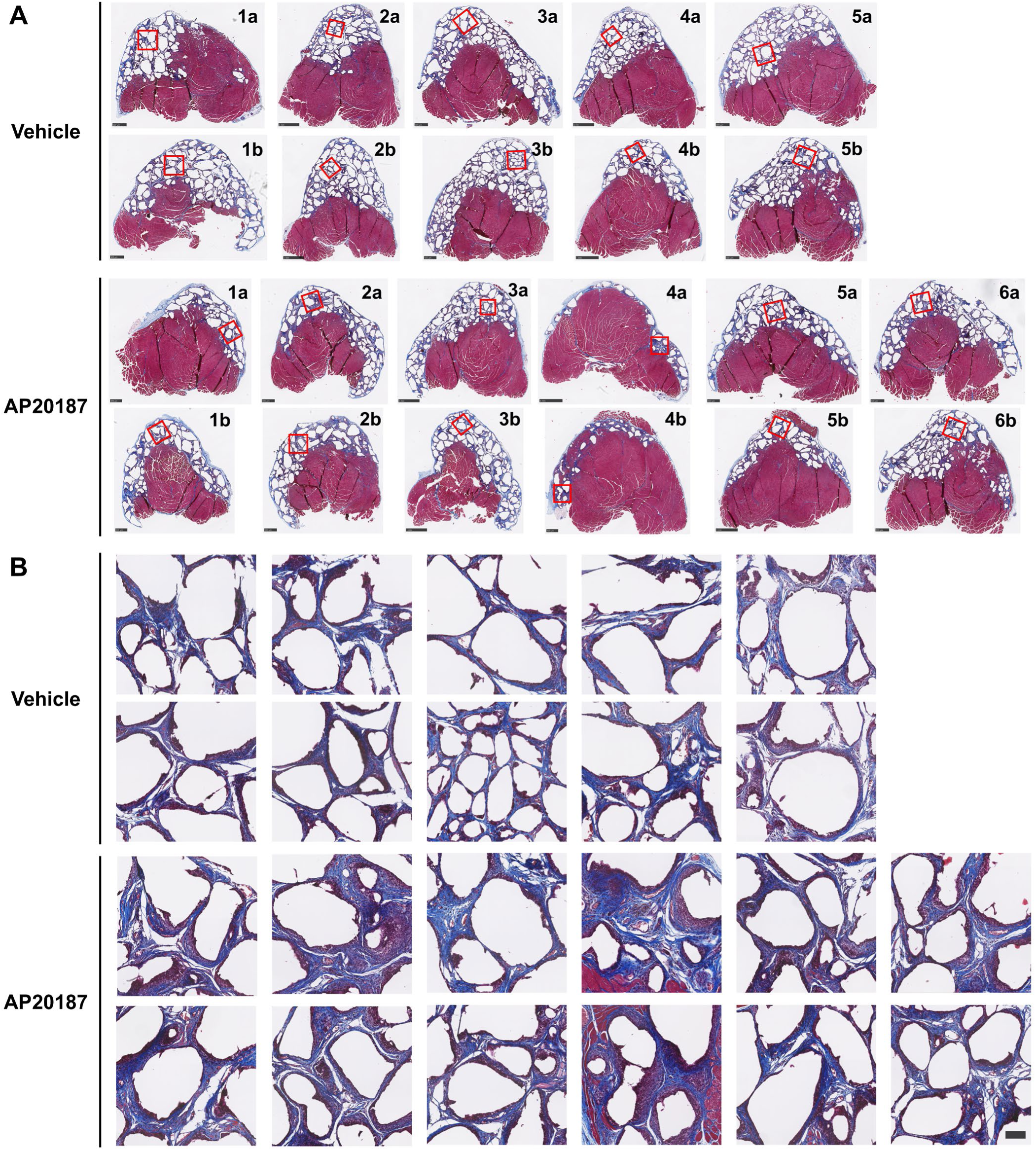
Fibrosis in implants with (AP20187) and without (vehicle) pericyte-lineage SnC elimination. **A)** Masson’s Trichrome staining of whole quadriceps implant tissue sections in each biological replicate (labeled by numbers). Two sections are shown per biological replicate (labeled by letters). **B)** Zoomed-in images of the regions outlined in red in **A**. Scale bars in **A**: 1000 µm for Vehicle 2a, 2b, 4a, 4b and AP20187 3a, 3b, 4a, 5b. 500 µm for Vehicle 1a, 1b, 3a, 3b, 5a, 5b and AP20187 1a, 1b, 2a, 2b, 4b, 5a, 6a, 6b. Scale bar in **B**: 100 µm

**Ext. Data 12.**
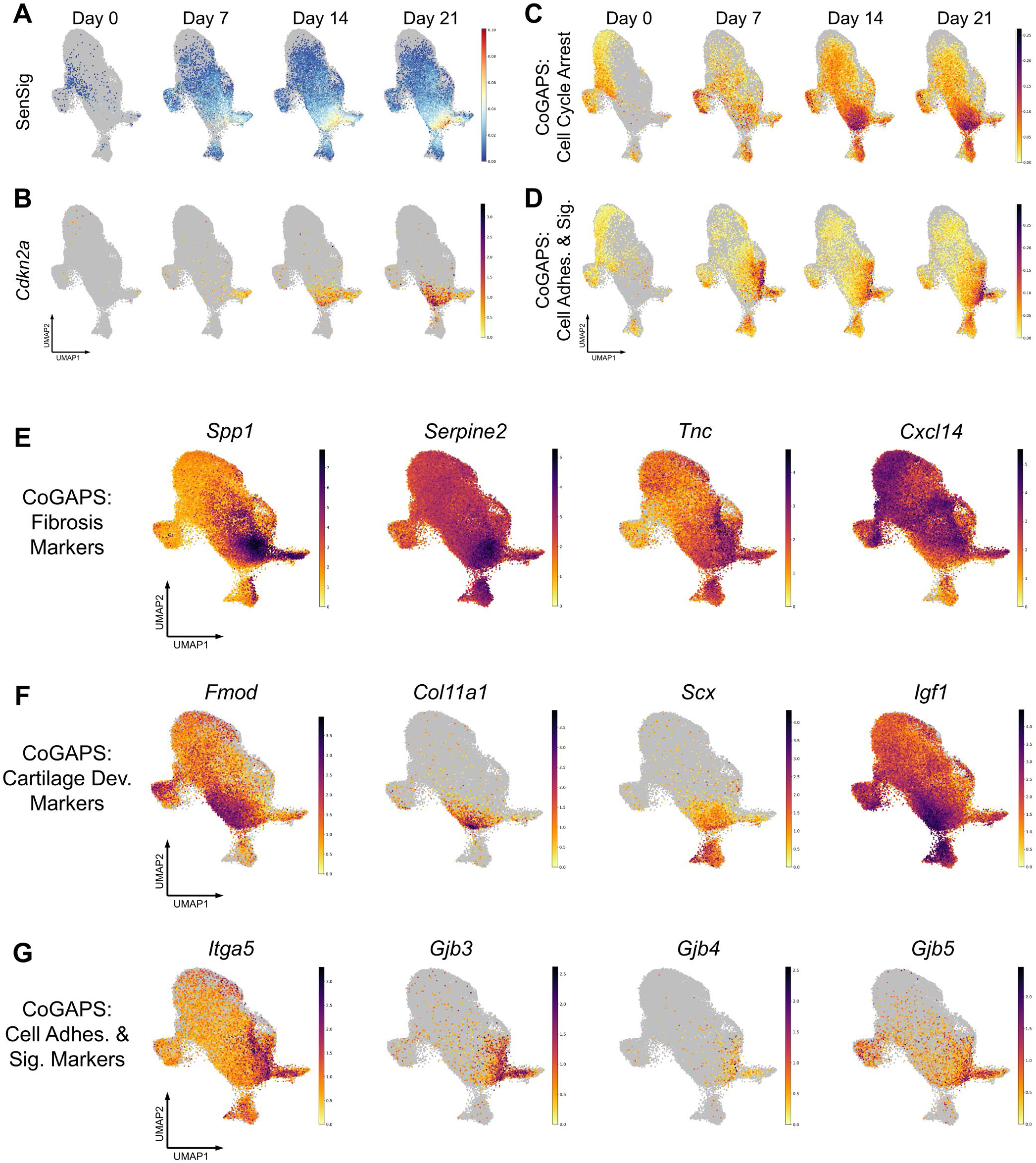
Expression of additional senotype patterns and markers in bleomycin-induced lung fibrosis. **A-D)** UMAP displaying SenSig projection weight (**A**), *Cdkn2a* normalized expression (**B**), CoGAPS Cell Cycle Arrest projection weight (**C**), and CoGAPS Cell Adhesion Projection weight (**D**) at Day 0 (no injury), 7, 14, and 21 following bleomycin-induced lung injury. **E-G)** UMAP displaying normalized expression of select marker genes from CoGAPS patterns Fibrosis (**E**), Cartilage Development (**F**), and Cell Adhesion & Signaling (**G**).

**Extended Data 13.**
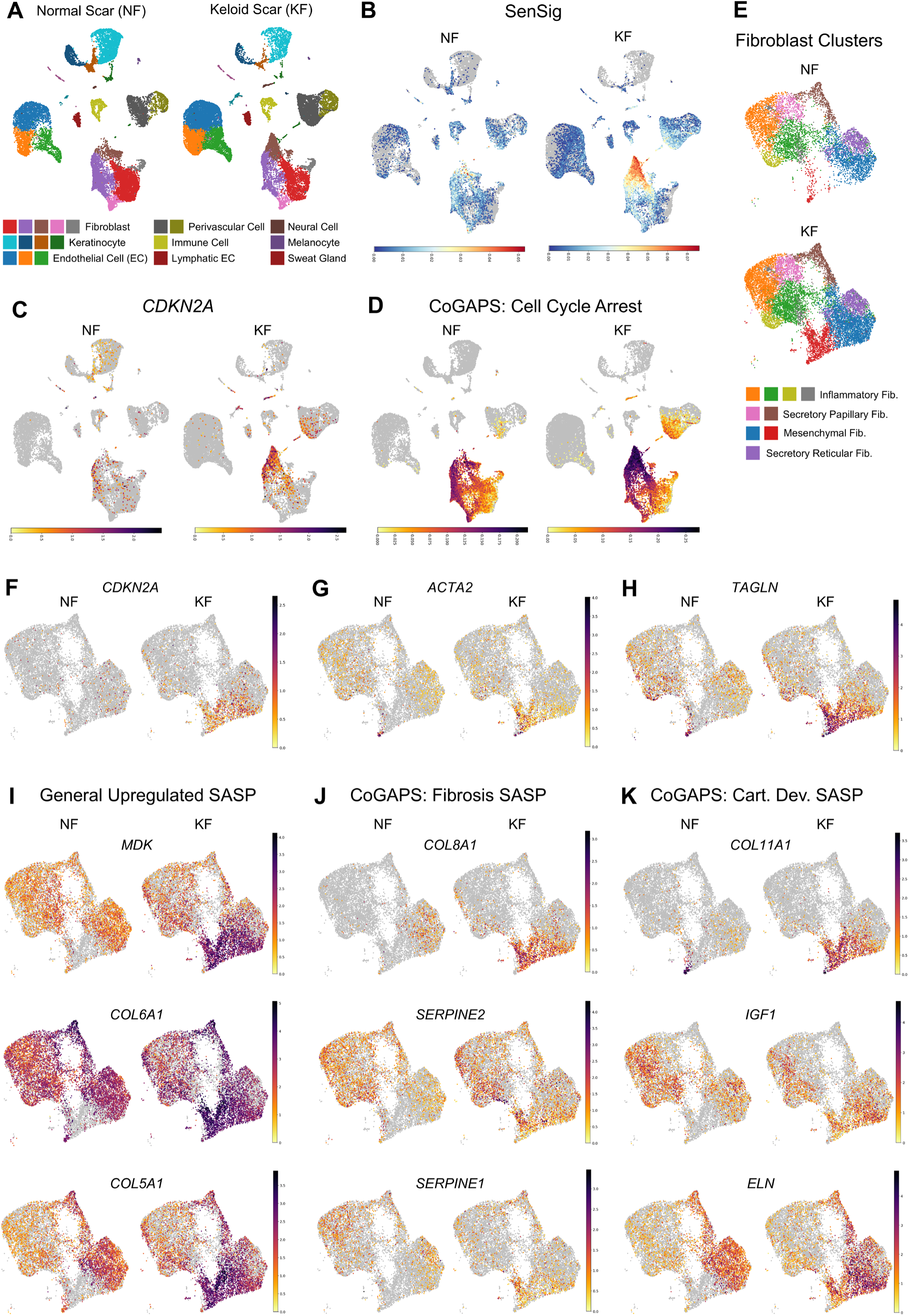
Expression of additional senotype markers in keloid scar. **A-D)** UMAP plots showing clusters (**A**), SenSig projection weight (**B**), *CDKN2A* normalized expression (**C**), and CoGAPS Cell Cycle Arrest projection weight (**D**) in all cell populations present in normal (NF) and keloid (KF) scars. **E)** Fibroblast subclusters in NF and KF. **F-H)** UMAP showing normalized expression of *CDKN2A* (**F**) and myofibroblast genes ACTA2 (**G**) and TAGLN (**H**) in fibroblast subclusters. **I)** Normalized expression of SASP factors upregulated across all SenSig+ cells in KF. **J-K)** Normalized expression of SASP genes from CoGAPS Fibrosis (**J**) and Cartilage Development (**K**) patterns that are upregulated in subpopulations of fibroblast SnCs in KF.

## References

1 Franzen, L. et al. Mapping spatially resolved transcriptomes in human and mouse pulmonary fibrosis. Nat Genet 56, 1725–1736 (2024). 10.1038/s41588-024-01819-2

2 Mayr, C. H. et al. Spatial transcriptomic characterization of pathologic niches in IPF. Sci Adv 10, eadl5473 (2024). 10.1126/sciadv.adl5473

3 Vannan, A. et al. Spatial transcriptomics identifies molecular niche dysregulation associated with distal lung remodeling in pulmonary fibrosis. Nat Genet 57, 647–658 (2025). 10.1038/s41588-025-02080-x

4 Rodriguez Morales, D., et al. Vascular Niches Are the Primary Hotspots in Cardiac Aging. Circ Res 137, 1353–1367 (2025). 10.1161/CIRCRESAHA.125.327060

5 Kuppe, C. et al. Decoding myofibroblast origins in human kidney fibrosis. Nature 589, 281–286 (2021). 10.1038/s41586-020-2941-1

6 Schafer, M. J. et al. Cellular senescence mediates fibrotic pulmonary disease. Nat Commun 8, 14532 (2017). 10.1038/ncomms14532

7 Mylonas, K. J. et al. Cellular senescence inhibits renal regeneration after injury in mice, with senolytic treatment promoting repair. Sci Transl Med 13 (2021). 10.1126/scitranslmed.abb0203

8 Belle, J. I. et al. Senescence Defines a Distinct Subset of Myofibroblasts That Orchestrates Immunosuppression in Pancreatic Cancer. Cancer Discov 14, 1324–1355 (2024). 10.1158/2159-8290.CD-23-0428

9 Yang, Y. et al. A BCL-xL/BCL-2 PROTAC effectively clears senescent cells in the liver and reduces MASH-driven hepatocellular carcinoma in mice. Nat Aging 5, 386–400 (2025). 10.1038/s43587-025-00811-7

10 Krizhanovsky, V. et al. Senescence of activated stellate cells limits liver fibrosis. Cell 134, 657–667 (2008). 10.1016/j.cell.2008.06.049

11 Meyer, K., Hodwin, B., Ramanujam, D., Engelhardt, S. & Sarikas, A. Essential Role for Premature Senescence of Myofibroblasts in Myocardial Fibrosis. J Am Coll Cardiol 67, 2018–2028 (2016). 10.1016/j.jacc.2016.02.047

12 Zhao, H. et al. Identifying specific functional roles for senescence across cell types. Cell 187, 7314–7334 e7321 (2024). 10.1016/j.cell.2024.09.021

13 O’Sullivan, E. D. et al. Single-cell analysis of senescent epithelia reveals targetable mechanisms promoting fibrosis. JCI Insight 7 (2022). 10.1172/jci.insight.154124

14 Demaria, M. et al. An essential role for senescent cells in optimal wound healing through secretion of PDGF-AA. Dev Cell 31, 722–733 (2014). 10.1016/j.devcel.2014.11.012

15 Jaiswal, A. et al. Spatial transcriptomics reveals altered communities and drivers of aberrant epithelia and pro-fibrotic fibroblasts in interstitial lung diseases. Cell Genom 6, 101066 (2026). 10.1016/j.xgen.2025.101066

16 Guo, J. L. et al. Histological signatures map anti-fibrotic factors in mouse and human lungs. Nature 641, 993–1004 (2025). 10.1038/s41586-025-08727-3

17 Bernier, L. P. et al. Brain pericytes and perivascular fibroblasts are stromal progenitors with dual functions in cerebrovascular regeneration after stroke. Nat Neurosci 28, 517–535 (2025). 10.1038/s41593-025-01872-y

18 Sato, R. et al. beta-III tubulin identifies anti-fibrotic state of pericytes in pulmonary fibrosis. Res Sq (2025). 10.21203/rs.3.rs-8138421/v1

19 Orvis, J. et al. gEAR: Gene Expression Analysis Resource portal for community-driven, multi-omic data exploration. Nat Methods 18, 843–844 (2021). 10.1038/s41592-021-01200-9

20 Ruta, A. et al. gammadelta T cell-stromal networks modulate matrix composition and vascularity in foreign body response. Nat Commun (2026). 10.1038/s41467-026-71540-7

21 Ruta, A. et al. gammadelta17 T cell-stromal networks modulate matrix composition and vascularity in foreign body response. bioRxiv (2025). 10.1101/2025.09.30.679608

22 Mejias, J. C. et al. Regulatory T cells clonally expand and contribute to stromal cell function in fibrotic response to synthetic implants. bioRxiv (2026). 10.64898/2026.01.05.697727

23 Sharma, G., Colantuoni, C., Goff, L. A., Fertig, E. J. & Stein-O’Brien, G. projectR: an R/Bioconductor package for transfer learning via PCA, NMF, correlation and clustering. Bioinformatics 36, 3592–3593 (2020). 10.1093/bioinformatics/btaa183

24 Stein-O’Brien, G. L. et al. Decomposing Cell Identity for Transfer Learning across Cellular Measurements, Platforms, Tissues, and Species. Cell Syst 8, 395–411 e398 (2019). 10.1016/j.cels.2019.04.004

25 Cherry, C. et al. Transfer learning in a biomaterial fibrosis model identifies in vivo senescence heterogeneity and contributions to vascularization and matrix production across species and diverse pathologies. Geroscience 45, 2559–2587 (2023). 10.1007/s11357-023-00785-7

26 Kowalczyk, M. S. et al. Single-cell RNA-seq reveals changes in cell cycle and differentiation programs upon aging of hematopoietic stem cells. Genome Res 25, 1860–1872 (2015). 10.1101/gr.192237.115

27 Hao, Y. et al. Dictionary learning for integrative, multimodal and scalable single-cell analysis. Nat Biotechnol 42, 293–304 (2024). 10.1038/s41587-023-01767-y

28 Victorelli, S. et al. Apoptotic stress causes mtDNA release during senescence and drives the SASP. Nature 622, 627–636 (2023). 10.1038/s41586-023-06621-4

29 Garcia-Alonso, L. et al. Single-cell roadmap of human gonadal development. Nature 607, 540–547 (2022). 10.1038/s41586-022-04918-4

30 Wang, R. et al. SEPDB: a database of secreted proteins. Database (Oxford) 2024 (2024). 10.1093/database/baae007

31 Shao, X., Taha, I. N., Clauser, K. R., Gao, Y. T. & Naba, A. MatrisomeDB: the ECM-protein knowledge database. Nucleic Acids Res 48, D1136–D1144 (2020). 10.1093/nar/gkz849

32 Bausch-Fluck, D. et al. The in silico human surfaceome. Proc Natl Acad Sci U S A 115, E10988–E10997 (2018). 10.1073/pnas.1808790115

33 Collins, B. C. et al. Multi-laboratory assessment of reproducibility, qualitative and quantitative performance of SWATH-mass spectrometry. Nat Commun 8, 291 (2017). 10.1038/s41467-017-00249-5

34 Gillet, L. C. et al. Targeted data extraction of the MS/MS spectra generated by data-independent acquisition: a new concept for consistent and accurate proteome analysis. Mol Cell Proteomics 11, O111 016717 (2012). 10.1074/mcp.O111.016717

35 Meier, F. et al. Online Parallel Accumulation-Serial Fragmentation (PASEF) with a Novel Trapped Ion Mobility Mass Spectrometer. Mol Cell Proteomics 17, 2534–2545 (2018). 10.1074/mcp.TIR118.000900

36 Moiseeva, V. et al. Senescence atlas reveals an aged-like inflamed niche that blunts muscle regeneration. Nature 613, 169–178 (2023). 10.1038/s41586-022-05535-x

37 Reyes, N. S. et al. Sentinel p16(INK4a+) cells in the basement membrane form a reparative niche in the lung. Science 378, 192–201 (2022). 10.1126/science.abf3326

38 Sturmlechner, I. et al. p21 produces a bioactive secretome that places stressed cells under immunosurveillance. Science 374, eabb3420 (2021). 10.1126/science.abb3420

39 Saul, D. et al. A new gene set identifies senescent cells and predicts senescence-associated pathways across tissues. Nat Commun 13, 4827 (2022). 10.1038/s41467-022-32552-1

40 Walter, L. D. et al. Transcriptomic analysis of skeletal muscle regeneration across mouse lifespan identifies altered stem cell states. Nat Aging 4, 1862–1881 (2024). 10.1038/s43587-024-00756-3

41 Stein-O’Brien, G. L. et al. Enter the Matrix: Factorization Uncovers Knowledge from Omics. Trends Genet 34, 790–805 (2018). 10.1016/j.tig.2018.07.003

42 Johnson, J. A. I. et al. Inferring cellular and molecular processes in single-cell data with non-negative matrix factorization using Python, R and GenePattern Notebook implementations of CoGAPS. Nat Protoc 18, 3690–3731 (2023). 10.1038/s41596-023-00892-x

43 Sherman, T. D., Gao, T. & Fertig, E. J. CoGAPS 3: Bayesian non-negative matrix factorization for single-cell analysis with asynchronous updates and sparse data structures. BMC Bioinformatics 21, 453 (2020). 10.1186/s12859-020-03796-9

44 Bidaut, G. & Ochs, M. F. ClutrFree: cluster tree visualization and interpretation. Bioinformatics 20, 2869–2871 (2004). 10.1093/bioinformatics/bth307

45 Brunet, J. P., Tamayo, P., Golub, T. R. & Mesirov, J. P. Metagenes and molecular pattern discovery using matrix factorization. Proc Natl Acad Sci U S A 101, 4164–4169 (2004). 10.1073/pnas.0308531101

46 Van de Sande, B. et al. A scalable SCENIC workflow for single-cell gene regulatory network analysis. Nat Protoc 15, 2247–2276 (2020). 10.1038/s41596-020-0336-2

47 Deshpande, A. et al. Uncovering the spatial landscape of molecular interactions within the tumor microenvironment through latent spaces. Cell Syst 14, 285–301 e284 (2023). 10.1016/j.cels.2023.03.004

48 Way, G. P., Zietz, M., Rubinetti, V., Himmelstein, D. S. & Greene, C. S. Compressing gene expression data using multiple latent space dimensionalities learns complementary biological representations. Genome Biol 21, 109 (2020). 10.1186/s13059-020-02021-3

49 Mohammadi, S., Davila-Velderrain, J. & Kellis, M. A multiresolution framework to characterize single-cell state landscapes. Nat Commun 11, 5399 (2020). 10.1038/s41467-020-18416-6

50 Chung, L. et al. Interleukin 17 and senescent cells regulate the foreign body response to synthetic material implants in mice and humans. Sci Transl Med 12 (2020). 10.1126/scitranslmed.aax3799

51 Tsukui, T., Wolters, P. J. & Sheppard, D. Alveolar fibroblast lineage orchestrates lung inflammation and fibrosis. Nature 631, 627–634 (2024). 10.1038/s41586-024-07660-1

52 Friscic, J. et al. The complement system drives local inflammatory tissue priming by metabolic reprogramming of synovial fibroblasts. Immunity 54, 1002–1021 e1010 (2021). 10.1016/j.immuni.2021.03.003

53 Correa-Gallegos, D. et al. CD201(+) fascia progenitors choreograph injury repair. Nature 623, 792–802 (2023). 10.1038/s41586-023-06725-x

54 Mascharak, S. et al. Multi-omic analysis reveals divergent molecular events in scarring and regenerative wound healing. Cell Stem Cell 29, 315–327 e316 (2022). 10.1016/j.stem.2021.12.011

55 Hernandez-Segura, A. et al. Unmasking Transcriptional Heterogeneity in Senescent Cells. Curr Biol 27, 2652–2660 e2654 (2017). 10.1016/j.cub.2017.07.033

56 Sadtler, K. et al. Developing a pro-regenerative biomaterial scaffold microenvironment requires T helper 2 cells. Science 352, 366–370 (2016). 10.1126/science.aad9272

57 Cable, D. M. et al. Robust decomposition of cell type mixtures in spatial transcriptomics. Nat Biotechnol 40, 517–526 (2022). 10.1038/s41587-021-00830-w

58 Palla, G. et al. Squidpy: a scalable framework for spatial omics analysis. Nat Methods 19, 171–178 (2022). 10.1038/s41592-021-01358-2

59 Alex, L. et al. Cardiac Pericytes Acquire a Fibrogenic Phenotype and Contribute to Vascular Maturation After Myocardial Infarction. Circulation 148, 882–898 (2023). 10.1161/CIRCULATIONAHA.123.064155

60 Quijada, P. et al. Cardiac pericytes mediate the remodeling response to myocardial infarction. J Clin Invest 133 (2023). 10.1172/JCI162188

61 Farr, J. N. et al. Local senolysis in aged mice only partially replicates the benefits of systemic senolysis. J Clin Invest 133 (2023). 10.1172/JCI162519

62 Suda, M. et al. Endothelial senescent-cell-specific clearance alleviates metabolic dysfunction in obese mice. Cell Metab 37, 2455–2465 e2456 (2025). 10.1016/j.cmet.2025.10.009

63 Deng, C. C. et al. Single-cell RNA-seq reveals fibroblast heterogeneity and increased mesenchymal fibroblasts in human fibrotic skin diseases. Nat Commun 12, 3709 (2021). 10.1038/s41467-021-24110-y

64 Darmawan, C. C. et al. Dasatinib Attenuates Fibrosis in Keloids by Decreasing Senescent Cell Burden. Acta Derm Venereol 103, adv4475 (2023). 10.2340/actadv.v103.4475

65 Kong, Y. X. et al. FOXO4-DRI induces keloid senescent fibroblast apoptosis by promoting nuclear exclusion of upregulated p53-serine 15 phosphorylation. Commun Biol 8, 299 (2025). 10.1038/s42003-025-07738-0

66 Limandjaja, G. C., Belien, J. M., Scheper, R. J., Niessen, F. B. & Gibbs, S. Hypertrophic and keloid scars fail to progress from the CD34(-) /alpha-smooth muscle actin (alpha-SMA)(+) immature scar phenotype and show gradient differences in alpha-SMA and p16 expression. Br J Dermatol 182, 974–986 (2020). 10.1111/bjd.18219

67 Basisty, N. et al. A proteomic atlas of senescence-associated secretomes for aging biomarker development. PLoS Biol 18, e3000599 (2020). 10.1371/journal.pbio.3000599

68 Schafer, M. J. et al. The senescence-associated secretome as an indicator of age and medical risk. JCI Insight 5 (2020). 10.1172/jci.insight.133668

69 Wang, B., Han, J., Elisseeff, J. H. & Demaria, M. The senescence-associated secretory phenotype and its physiological and pathological implications. Nat Rev Mol Cell Biol 25, 958–978 (2024). 10.1038/s41580-024-00727-x

70 Ye, J. et al. Senescent CAFs Mediate Immunosuppression and Drive Breast Cancer Progression. Cancer Discov 14, 1302–1323 (2024). 10.1158/2159-8290.CD-23-0426

71 Schilling, B. et al. Senescence-Linked Fibrosis in the Aging Human Ovary Revealed by p16-Based Histological Profiling and Spatial Transcriptomics. Res Sq (2026). 10.21203/rs.3.rs-8290960/v1

72 Saul, D. et al. Osteochondroprogenitor cells and neutrophils expressing p21 and senescence markers modulate fracture repair. J Clin Invest 134 (2024). 10.1172/JCI179834

73 Gasek, N. S. et al. Clearance of p21 highly expressing senescent cells accelerates cutaneous wound healing. Nat Aging 5, 21–27 (2025). 10.1038/s43587-024-00755-4

74 Wang, B. et al. Intermittent clearance of p21-highly-expressing cells extends lifespan and confers sustained benefits to health and physical function. Cell Metab 36, 1795–1805 e1796 (2024). 10.1016/j.cmet.2024.07.006

75 Meguro, S. et al. Preexisting senescent fibroblasts in the aged bladder create a tumor-permissive niche through CXCL12 secretion. Nat Aging 4, 1582–1597 (2024). 10.1038/s43587-024-00704-1

76 Chambers, E. S. et al. Recruitment of inflammatory monocytes by senescent fibroblasts inhibits antigen-specific tissue immunity during human aging. Nat Aging 1, 101–113 (2021). 10.1038/s43587-020-00010-6

77 Assouline, B. et al. Senescent cancer-associated fibroblasts in pancreatic adenocarcinoma restrict CD8(+) T cell activation and limit responsiveness to immunotherapy in mice. Nat Commun 15, 6162 (2024). 10.1038/s41467-024-50441-7

78 Liu, Y. et al. Conserved spatial subtypes and cellular neighborhoods of cancer-associated fibroblasts revealed by single-cell spatial multi-omics. Cancer Cell 43, 905–924 e906 (2025). 10.1016/j.ccell.2025.03.004

79 Mebane, R. H. et al. Spatial transcriptomic analysis of immune checkpoint blockade response in triple negative breast cancers with tertiary lymphoid structures. iScience 28, 112808 (2025). 10.1016/j.isci.2025.112808

80 Acosta, J. C. et al. A complex secretory program orchestrated by the inflammasome controls paracrine senescence. Nat Cell Biol 15, 978–990 (2013). 10.1038/ncb2784

81 Hubackova, S., Krejcikova, K., Bartek, J. & Hodny, Z. IL1- and TGFbeta-Nox4 signaling, oxidative stress and DNA damage response are shared features of replicative, oncogene-induced, and drug-induced paracrine ’bystander senescence’. Aging (Albany NY) 4, 932–951 (2012). 10.18632/aging.100520

82 Hoare, M. et al. NOTCH1 mediates a switch between two distinct secretomes during senescence. Nat Cell Biol 18, 979–992 (2016). 10.1038/ncb3397

83 Bird, T. G. et al. TGFbeta inhibition restores a regenerative response in acute liver injury by suppressing paracrine senescence. Sci Transl Med 10 (2018). 10.1126/scitranslmed.aan1230

84 Teo, Y. V. et al. Notch Signaling Mediates Secondary Senescence. Cell Rep 27, 997–1007 e1005 (2019). 10.1016/j.celrep.2019.03.104

85 Vizovisek, M., Fonovic, M. & Turk, B. Cysteine cathepsins in extracellular matrix remodeling: Extracellular matrix degradation and beyond. Matrix Biol 75–76, 141–159 (2019). 10.1016/j.matbio.2018.01.024

86 Biel, C., Faber, K. N., Bank, R. A. & Olinga, P. Matrix metalloproteinases in intestinal fibrosis. J Crohns Colitis 18, 462–478 (2024). 10.1093/ecco-jcc/jjad178

87 van Splunder, H., Villacampa, P., Martinez-Romero, A. & Graupera, M. Pericytes in the disease spotlight. Trends Cell Biol 34, 58–71 (2024). 10.1016/j.tcb.2023.06.001

88 Dias, D. O. et al. Pericyte-derived fibrotic scarring is conserved across diverse central nervous system lesions. Nat Commun 12, 5501 (2021). 10.1038/s41467-021-25585-5

89 Zhu, X. et al. Age-dependent fate and lineage restriction of single NG2 cells. Development 138, 745–753 (2011). 10.1242/dev.047951

90 O’Flanagan, C. H. et al. Dissociation of solid tumor tissues with cold active protease for single-cell RNA-seq minimizes conserved collagenase-associated stress responses. Genome Biol 20, 210 (2019). 10.1186/s13059-019-1830-0

91 Chen, Y., Chen, L., Lun, A. T. L., Baldoni, P. L. & Smyth, G. K. edgeR v4: powerful differential analysis of sequencing data with expanded functionality and improved support for small counts and larger datasets. Nucleic Acids Res 53 (2025). 10.1093/nar/gkaf018

92 Liberzon, A. et al. The Molecular Signatures Database (MSigDB) hallmark gene set collection. Cell Syst 1, 417–425 (2015). 10.1016/j.cels.2015.12.004

93 Milacic, M. et al. The Reactome Pathway Knowledgebase 2024. Nucleic Acids Res 52, D672–D678 (2024). 10.1093/nar/gkad1025

94 Ashburner, M. et al. Gene ontology: tool for the unification of biology. The Gene Ontology Consortium. Nat Genet 25, 25–29 (2000). 10.1038/75556

95 Gene Ontology, C. The Gene Ontology knowledgebase in 2026. Nucleic Acids Res 54, D1779–D1792 (2026). 10.1093/nar/gkaf1292

96 Castanza, A. S. et al. Extending support for mouse data in the Molecular Signatures Database (MSigDB). Nat Methods 20, 1619–1620 (2023). 10.1038/s41592-023-02014-7

97 Lvovs, D. et al. STAPLE: automating spatial transcriptomics analysis and AI interpretation. bioRxiv (2026). 10.64898/2026.03.30.715127

98 Virshup, I., Rybakov, S., Theis, F. J., Angerer, P. & Wolf, F. A. anndata: Access and store annotated datamatrices. Journal of Open Source Software 9 (2024). 10.21105/joss.04371

99 Wickham, H. *ggplot2: Elegant Graphics for Data Analysis*. (Springer-Verlag New York, 2016).

100 Wickham H, A. M., Bryan J, Chang W, McGowan LD, François R, Grolemund G, Hayes A, Henry L, Hester J, Kuhn M, Pedersen TL, Miller E, Bache SM, Müller K, Ooms J, Robinson D, Seidel DP, Spinu V, Takahashi K, Vaughan D, Wilke C, Woo K, Yutani H. Welcome to the tidyverse. Journal of Open Source Software 4, 1686 (2019). 10.21105/joss.01686

101 Gu, Z. Complex heatmap visualization. Imeta 1, e43 (2022). 10.1002/imt2.43

102 Gu, Z., Eils, R. & Schlesner, M. Complex heatmaps reveal patterns and correlations in multidimensional genomic data. Bioinformatics 32, 2847–2849 (2016). 10.1093/bioinformatics/btw313

